# Evolutionarily conserved roles for blood-brain barrier xenobiotic transporters in endogenous steroid partitioning and behavior

**DOI:** 10.1101/131698

**Authors:** Samantha J. Hindle, Roeben N. Munji, Elena Dolghih, Garrett Gaskins, Souvinh Orng, Hiroshi Ishimoto, Allison Soung, Michael DeSalvo, Toshihiro Kitamoto, Michael J. Keiser, Matthew P. Jacobson, Richard Daneman, Roland J. Bainton

## Abstract

Optimal brain function depends upon efficient control over the brain entry of blood components; this is provided by the blood-brain barrier (BBB). Curiously, some brain-impermeable drugs can still cause behavioral side effects.

To investigate this phenomenon, we asked whether the promiscuous drug efflux transporter Mdr1 has dual functions in transporting drugs and endogenous molecules. If this is true, brain-impermeable drugs may cause behavioral side effects by affecting brain levels of endogenous molecules.

Using computational, genetic and pharmacologic approaches across diverse organisms we demonstrate that BBB-localized efflux transporters are critical for regulating brain levels of endogenous steroids, and steroid-regulated behaviors (sleep in *Drosophila* and anxiety in mice). Furthermore, we show that Mdr1-interacting drugs are associated with anxiety-related behaviors in humans.

We propose a general mechanism for common behavioral side effects of prescription drugs: pharmacologically challenging BBB efflux transporters disrupts brain levels of endogenous substrates, and implicates the BBB in behavioral regulation.

#### Abbreviations

20-E: 20-hydroxyecdysone
ABC: ATP-binding Cassette
ADRs: Adverse Drug Reactions
BBB: Blood-Brain Barrier
BVEC: brain vascular endothelial cells
CNS: central nervous system
EcRLBD: ecdysone receptor ligand binding domain
EF: enrichment factor
MDR: multidrug resistant
RhoB: Rhodamine B
SPG: Subperineurial glia

## Introduction

The maintenance of central nervous system (CNS) function depends upon its regulated isolation from circulating drugs and toxins (xenobiotics), as well as endogenous molecules produced in the periphery. The importance of insulating neural tissue is evidenced by the conserved molecular and anatomic properties of blood-brain barriers (BBBs) present throughout evolutionarily diverse organisms (Fig. 1A). The vertebrate BBB is composed of brain vascular endothelial cells (BVECs), pericytes, basal lamina, and the endfeet of astrocytes. BVECs are specialized to protect the CNS as, compared to peripheral endothelial cells, they are enriched for tight junction components, chemoprotective ATP-Binding Cassette (ABC) drug efflux transporters and nutrient influx transporters (Daneman et al., 2010, Daneman, 2012). Similar to vertebrate BVECs, *Drosophila melanogaster* focuses chemoprotective physiology into a single functionally polarized cellular layer, the subperineurial glia (SPG). The SPG layer surrounds the CNS, separating it from the blood (hemolymph) (Stork et al., 2008), and similarly possesses the hallmark properties of the vertebrate BBB: tight junctional physiology, potent xenobiotic efflux biology and particular metabolic transport characteristics (Mayer et al., 2009, DeSalvo et al., 2014, Hindle and Bainton, 2014).

**Figure 1.**
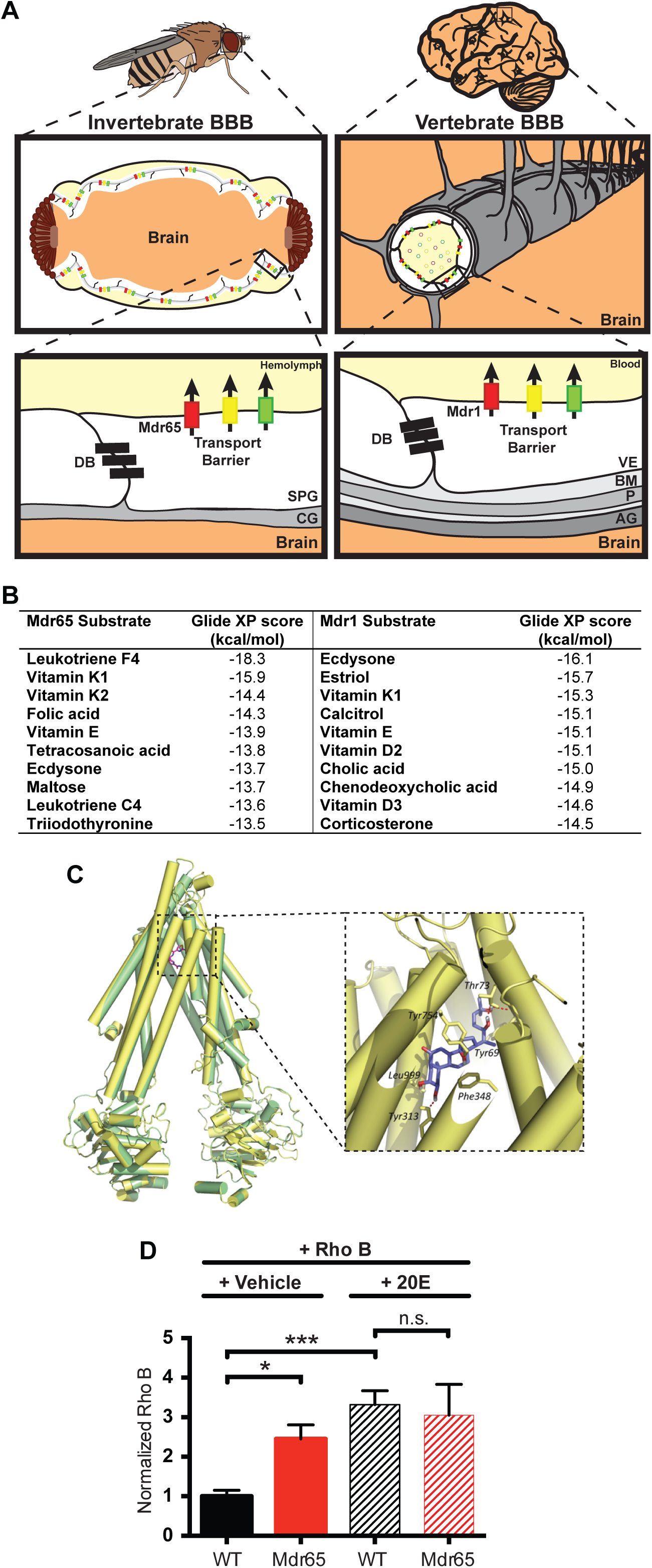
The *Drosophila* steroid hormone 20-hydroxyecdysone is a predicted Mdr65 substrate. (A) Diagramatical representation of the blood-brain barrier in invertebrates and vertebrates. The invertebrate BBB is a compound structure, consisting of the subperineurial glia (SPG), the outer perineurial glia, and a basement membrane known as the neural lamella (only the SPG layer is shown for simplicity). The vertebrate BBB is primarily formed by the vascular endothelial cells (VE) that form the capillaries in the brain, and its functions are supported by the surrounding pericytes (P) within the basement membrane (BM), and the end feet of the astrocyte glia (AG). These cellular and non-cellular layers form a compound barrier structure known as the neurovascular unit (NVU). Both the vertebrate and invertebrate barriers express junctional proteins that make up the diffusion barrier (DB), as well as various ATP-binding Cassette (ABC) transporters that form the transport barrier and protect the brain from xenobiotics. CG; cortex glia. (B) Substrate predictions for the fly Mdr65 and mouse Mdr1 efflux transporters from substrate docking computer modeling. (C) Homology model of Mdr65 (yellow) overlaid on the mouse Mdr1 template PDB ID: 3G60 (green). Original template ligand, QZ59-RRR, is shown in pink. The enlarged window shows the top-scored pose of ecdysone docked in the Mdr65 model. Also shown are some of the residues predicted to be involved in hydrogen bonding and hydrophobic interactions with the ligand. (D) Competitive *in vivo* efflux transport assay using Rhodamine B and 20-E. Fluorescence readings are normalized to control brains injected with Rhodamine B and vehicle. At least 4 biological replicates were performed for each condition. Error bars represent the SEM. ANOVA *, p<0.05; ***, p≤0.001.

Peripherally produced hydrophilic molecules, like catecholamines, are physically isolated from the CNS due to the BVEC tight junctions; this crucially allows synaptic regulation that is independent from the peripheral nervous system (Tsukita et al., 2001). Small lipophilic molecules, however, are able to freely diffuse across the plasma membrane, requiring an active chemical transport barrier to limit their buildup in the brain. Due to its lumenal membrane localization, ABC Type B efflux transporters, like vertebrate Mdr1, can provide this function, using ATP to efficiently prevent the CNS accumulation of its substrates (e.g. many lipophilic drugs) (Loscher and Potschka, 2005b, van Asperen et al., 1996, Loscher and Potschka, 2005a).

Mdr1 (P-glycoprotein/ABCB1) is expressed at the BBB and peripherally, in sites such as gut, kidney and other endothelial cells (Thiebaut et al., 1987, Cordon-Cardo et al., 1989). Unlike most transporters, it has a very broad spectrum of substrates, including calcium channel blockers, calmodulin antagonists, cyclic peptides, and endogenous molecules such as vitamins and steroids (Tsuruo et al., 1982a, Tsuruo et al., 1982b, Tsuruo et al., 1983, Naito et al., 1989, Silbermann et al., 1989, Ueda et al., 1992, Ford, 1996, Uhr et al., 2002, Muller et al., 2003, Schoenfelder et al., 2012). Thus, due to its promiscuous nature and physiologic localizations, Mdr1 is likely to have pharmacological interactions with many endogenous molecules, some of which have direct effects on the CNS. Indeed, acute, exogenous dosing of progesterone, aldosterone, cortisol or corticosterone in Mdr1 loss-of-function mice (Mdr1 KO) resulted in higher CNS levels (Uhr et al., 2002). Thus, the chemoprotective role of Mdr1 could have consequences on steady state brain disposition of potent endobiotics. Curiously, previous studies on Mdr1 KO animals that showed increased emotional stress behaviors, highlighted a peripheral endobiotic role for Mdr1 (Schoenfelder et al, 2012). However, a specific BBB role for Mdr1 that links endobiotic clearance mechanisms and behavioral regulation has not yet been demonstrated. To address this we took a broad look at potential endobiotic substrates and BBB-enriched ABC transporters to specifically investigate their BBB role in brain steroid regulation.

We first used a *Drosophila* model of xenobiotic efflux transporter deficiency to investigate its conserved roles in CNS disposition of endogenous molecules and in discrete animal behaviors. We previously identified the *Drosophila* Mdr1 homolog (Mdr65) and showed it is highly enriched at the BBB, functions as a chemoprotective efflux transporter, and can transport vertebrate Mdr1 substrates (DeSalvo et al., 2014, Mayer et al., 2009). Here, we advance these findings by using computational and genetic studies in *Drosophila,* followed by mass spectrometry on mouse CNS to show evolutionarily-conserved roles for BBB ABC transporters in regulating levels of CNS endobiotics (i.e. ecdysone in *Drosophila* and aldosterone in mice). Furthermore, xenobiotic transporter deficiency coincides with the dysregulation of a host of steroid-controlled physiologies, including developmental timing and sleep in *Drosophila* and anxiety in mice. We also show that acute, pharmacological inhibition of Mdr1 causes an accumulation of aldosterone in the mouse brain, and a set of similar Mdr1-interacting drugs/substrates are also significantly associated with CNS-related disorders (e.g. anxiety) in humans.

Together these findings suggest an important, evolutionarily-conserved role for BBB-localized chemoprotective ABC transporters that reaches beyond xenobiotics into endobiotic partitioning and behavioral regulation. Moreover, these data suggest that a common side effect of many drugs may have a general mechanism of action that depends, in part, on the altered CNS homeostasis of endogenous signaling molecules. The ability of the BBB to sense and respond to xenobiotics as well as changes in endogenous molecule partitioning provides a mechanism for how the BBB can efficiently sense and respond to chemical insult.

## Results

### Substrate docking and competitive transport assays suggest steroid hormones as substrates of xenobiotic efflux transporters

To first determine whether *Drosophila* is a suitable model for investigating endogenous substrate partitioning roles of Mdr1-like transporters, we took an unbiased approach to compare the potential substrates of the mouse Mdr1 and the *Drosophila* homologue Mdr65. Following prior work on human Mdr1 (Dolghih et al., 2011), we created a homology model of Mdr65 from a mouse Mdr1 template structure and used induced fit docking to virtually screen over 300 endogenous small molecules from the KEGG database. For both Mdr1 and Mdr65, docking predicted various endogenous molecules with potent biological functions, including hormones and vitamins, as potential substrates (Fig. 1B). As previously suggested (Uhr et al., 2002, Schoenfelder et al., 2012), we found that steroids are predicted substrates of mammalian Mdr1. Interestingly, an invertebrate steroid (ecdysone) was also a predicted substrate of the *Drosophila* Mdr65 transporter, which suggests that regulation of both xenobiotics and endogenous steroids is a conserved, and therefore important, role for Mdr1-like transporters.

To determine *in vivo* whether steroids are substrates of Mdr65, we performed a competitive efflux transport assay using the active form of ecdysone, 20-hydroxyecdysone, (20-E) and a known fluorescent Mdr65 substrate, Rhodamine B (RhoB) (Mayer et al., 2009). We tested two conditions: first we loaded brains with RhoB by co-injecting the hemolymph (fly blood) with RhoB (200 ng) and vehicle (20% ethanol), second we co-injected 200 ng RhoB and approximately 500 ng 20E (in 20% ethanol). After a 2-hour recovery period, we assessed the amount of fluorescence remaining in wild type and Mdr65 mutant brains. As expected, after RhoB and vehicle injection, Mdr65 efficiently removed RhoB from wild type brains, but in its absence, RhoB remained in the brain. Interestingly, after co-injection of RhoB and 20-E, we observed a 3-fold increase in fluorescence in wild type brains compared to co-injection of RhoB and vehicle (Fig. 1D). Furthermore, under the same conditions, there was no significant additional increase in fluorescence in Mdr65 mutant brains. These data show that 20E can compete with RhoB for its removal by Mdr65, and can function as a substrate of Mdr65.

### Blood-brain barrier partitioning of 20-hydroxyecdysone is altered in Mdr65 mutant flies

We next investigated whether 20E partitioning between the hemolymph and CNS was disrupted in Mdr65 mutants. We used an ecdysone reporter fly line (EcRLBD>Stinger GFP) to visualize the presence of ecdysone in the CNS. This reporter relies on a fusion protein of the Ecdysone receptor (EcR) ligand binding domain (LBD) and the Gal4 transcription factor (Palanker et al., 2006). Upon ecdysone binding, the fusion protein translocates to the nucleus and activates transcription of nuclear-localized GFP (Stinger GFP). We first assessed the time-course of induction of GFP after injecting approximately 500 ng 20E into the hemolymph of the fly. GFP was efficiently induced 16 hours following 20E injection and no induction was seen after vehicle injection (Fig. 2A).

**Figure 2.**
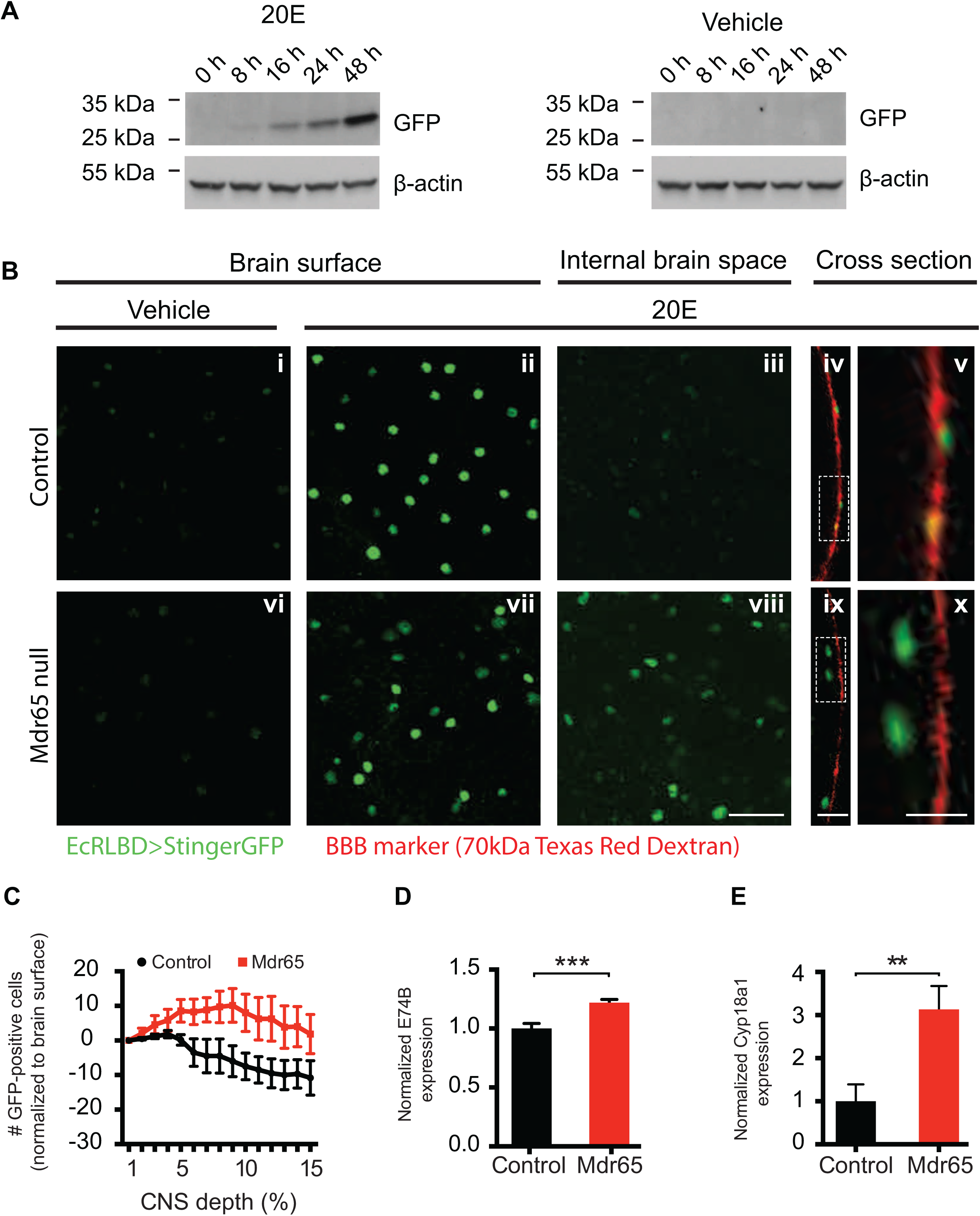
*Drosophila* Mdr65 null mutants show altered blood-brain barrier partitioning of 20-hydroxyecdysone. (A) Western blot time-course of EcRLBD>Stinger GFP fly heads following injection of 500 ng 20-E or vehicle (20% ethanol). The levels of β-actin were used as a loading control. N=2 technical replicates. (B) Confocal images of wild type (i-v) and Mdr65 null mutant (vi-x) brains expressing EcRLBD>Stinger GFP 16-24h after hemolymph injection of vehicle or approximately 500 ng 20E. The flies were co-injected with 70kDa Texas Red Dextran to mark the BBB. Images are shown at the level of the BBB layer (Brain surface; i, ii, vi & vii), below the BBB layer (Internal brain space; iii & viii) and at the cross section of the optic lobe (Cross section; iv, v, ix & x). Regions from the cross sections were enlarged for clarity (v & x). N> 9 biological replicates. Scale bars, 20 μm (i-iii, vi-viii), 10 μm (iv & ix) and 5 μm (v & x). (C) Quantification of the total number of GFP-positive cells/z-stack section for the initial 15% depth into the optic lobes of EcRLBD>Stinger GFP wild type and Mdr65 null flies hemolymph-injected 16-24h prior with approximately 500 ng 20E. Data were normalized to the number of GFP-positive cells present on the surface of each optic lobe. N > 5 biological replicates. (D & E) QPCR analysis of *E74B* (D) and *Cyp18a1* (E) transcript levels in whole brains from wild type and Mdr65 null flies (without the EcRLBD>Stinger GFP reporter). Error bars represent the SEM. T test **, p<0.005. See also Figures S3-5.

We then assessed whether 20E levels were increased in whole brain mounts from Mdr65 null mutants. In both control and Mdr65 mutant flies, hemolymph injection of 20E caused an increase in GFP fluorescence at the external surface of the brain, compared to injection of vehicle alone (Fig. 2B). However, GFP-positive cells were barely visible inside the brains of control flies. These data show that the reporter successfully indicates the presence of ecdysone in whole brain mounts, and that the wild type BBB is able to restrict 20E access to the brain. In contrast to the wild type BBB, loss of Mdr65 function resulted in an increased induction of the ecdysone reporter inside Mdr65 null mutant brains, as shown by single-plane images below the BBB (internal brain space) and at the brain cross section (Fig. 2B). The number of GFP-positive cells inside the brain was also quantified for each optical section of the z-stack series (Fig. 2C). These data clearly show an increased number of GFP-positive cells, and therefore increased levels of ecdysone, inside the brains of Mdr65 null mutants.

To further assess whether ecdysone levels were increased in Mdr65 mutant brains, we determined whether downstream ecdysone responses were altered. Ecdysone binds to the EcR nuclear hormone receptor, which dimerizes with the Retinoid-X receptor (RXR) homologue Ultraspiracle, and drives expression of ecdysone–responsive genes including *E74B* and *Cyp18a1* (Burtis et al., 1990, Thummel et al., 1990, Segraves and Hogness, 1990, Koelle et al., 1991, Hurban and Thummel, 1993). We found that the whole brain transcript levels of *E74B* were significantly higher in Mdr65 null brains compared to wild type brains (Fig. 2D). We also revealed metabolic consequences of diminished CNS chemoprotection as the 20E-inducible Cytochrome P450 *Cyp18a1* was increased 3-fold in Mdr65 mutant brains (Fig. 2E).

### Mdr1 regulates the brain levels of Aldosterone in mice

Our findings in the *Drosophila* system and Mdr1 substrate docking predictions suggest that BBB-enriched Mdr1, the closest mouse homolog to Mdr65, might also function to inhibit blood-to-CNS flux of circulating endogenous steroids in vertebrates. As the diversity of steroids in mammals is much more sophisticated than in *Drosophila,* we took a broad and unbiased HPLC-MS/MS approach to profile the CNS levels of steroid hormones in Mdr1 mutant mice. Mice have two Mdr1 genes, Mdr1a and Mdr1b. Thus we compared the brain levels of steroids in Mdr1 double knockout mice to control mice. Our analyses show that deletion of Mdr1 does not lead to statistically significant changes in the brain levels of endogenous androstenedione, androsterone, corticosterone, dihydrotestosterone (DHT), estrone, progesterone or testosterone. However, the brains of Mdr1 mutant mice have significantly higher levels of endogenous aldosterone than brains of control mice (Aldosterone, mean of raw values: control 0.088 nM, Mdr1: 0.132 nM; relative amount (normalized to mean of control): control 100, Mdr1: 168) (Fig. 3A). This result shows that Mdr1 function is necessary for maintaining normal brain levels of aldosterone in mice.

**Figure 3.**
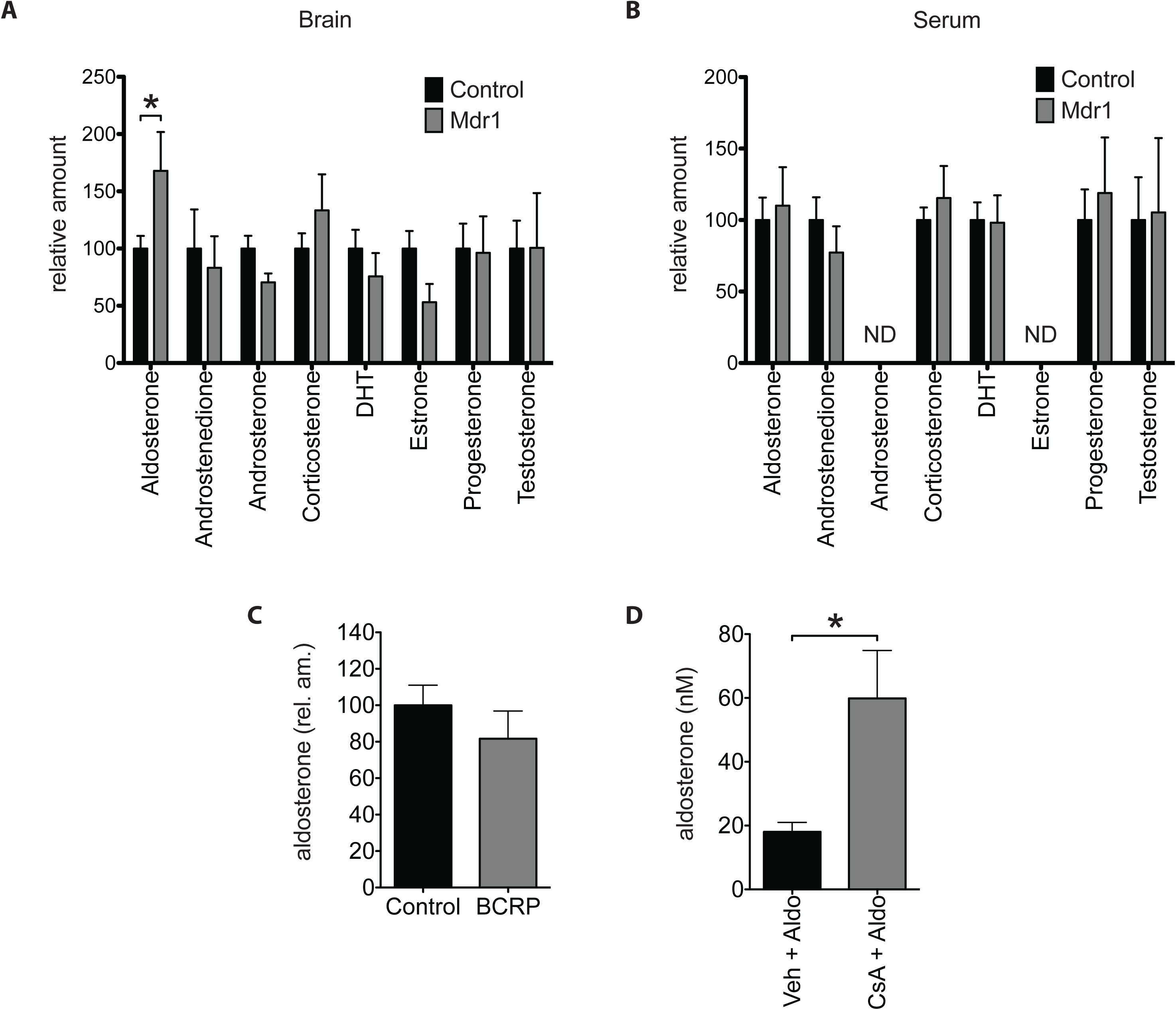
Endogenous aldosterone levels are increased in Mdr1 mutant mice. (A and B) HPLC-MSMS analysis of endogenous steroids from whole brains (A) and sera (B) of adult male Mdr1 control and mutant mice. N= 18 control and 7 Mdr1 mutant mice. ND, no data. (C) HPLC-MSMS analysis of endogenous aldosterone from whole brains of adult male BCRP control and mutant mice. N=18 Controls and 9 BCRP mutant mice. (D) HPLC-MSMS analysis of aldosterone from whole brains of wild type mice injected first with either vehicle (n=4) or Cyclosporin A (CsA) (n=4) then followed by aldosterone. Error bars represent SEM. Mann-Whitney test * p<0.05.

Aldosterone is synthesized and secreted into the blood by the adrenal cortex of the adrenal gland (Bollag, 2014). Our observation that deletion of Mdr1 leads to increased brain aldosterone suggests that there is an increase in blood-to-brain flux of aldosterone in Mdr1 mutant mice. However, this increase in blood-to-brain flux of aldosterone could also be attributed to the steepening of the blood-to-brain gradient of aldosterone caused by an increase in blood aldosterone. To verify that elevated aldosterone in Mdr1 mutant brains is not due to an increase in blood aldosterone, we measured steroid levels in serum as well. In contrast to brain, the serum levels of aldosterone and other steroids remained similar between Mdr1 control and mutant mice (Aldosterone, mean of raw values: control 0.573 nM, Mdr1: 0.657 nM; relative amount (normalized to mean of control): control: 100, Mdr1: 110) (Fig. 3B). These findings suggest that a steepening of the blood-to-brain gradient of aldosterone does not contribute to the observed elevation in brain aldosterone of Mdr1 mutant mice.

To further assess whether blood-to-brain flux of aldosterone is regulated by Mdr1, we tested whether acute pharmacological inhibition of Mdr1 can increase accumulation of aldosterone in the brain. For this experiment we used the established Mdr1 competitive inhibitor cyclosporin A (CsA) (Ejendal and Hrycyna, 2005, Elsinga et al., 2005) (Bakhsheshian et al., 2013, Miller, 2010). A caveat of using CsA is that CsA has also been shown to inhibit the BBB-enriched, efflux ABC transporter BCRP/ABCG2 (Bakhsheshian et al., 2013, Miller, 2010). Thus, we first established whether BCRP regulates the brain levels of aldosterone. As we did for Mdr1, we compared the levels of aldosterone in brains of control and BCRP mutant mice by HPLC-MS/MS. We determined that the brain levels of aldosterone are similar between control and BCRP mutant mice (Aldosterone, mean of raw values: control: 0.088 nM, BCRP: 0.064 nM; relative amount (normalized to mean of control), control: 100, BCRP: 82) (Fig. 3C). This result shows that the brain levels of aldosterone are not regulated by BCRP. Next, we proceeded to test whether acute inhibition of Mdr1 can increase accumulation of aldosterone in the brain. First we intraperitoneally injected adult male wild type mice with either the Mdr1 inhibitor CsA (50 mg/Kg bw) or vehicle. One hour later, we injected all mice with aldosterone (14 mg/Kg bw) intravenously by tail vein. Three hours post aldosterone injection, we harvested brains and measured brain aldosterone by HPLC-MS/MS. Brains of mice treated with CsA accumulated 3.3-fold more aldosterone than brains of mice treated with vehicle (vehicle: 18.032 nM, CsA: 59.872 nM) (Fig. 3D). This result shows that acute inhibition of Mdr1 with CsA increases blood-to-brain flux of aldosterone. Taken together, our results suggest that BBB Mdr1 regulates the blood-to-brain flux of aldosterone in mice.

### Blood-brain barrier Mdr65 regulates *Drosophila* behavior

As our experiments established a conserved function for xenobiotic ABC transporters in regulating endogenous steroids in flies and mice, we next investigated whether they also have conserved functions in regulating animal behavior. First, we assessed whether ecdysone-regulated behaviors were perturbed in flies deficient for Mdr65. It is well established that the behavioral transitions required for properly timed ecdysis and eclosion are dependent upon the precise timing of 20-hydroxyecdysone (20E) steroid peaks (Truman, 1981, Curtis et al., 1984, Truman, 1996, Zitnan et al., 1999, Zitnan et al., 2007). These behavioral transitions are also regulated, in part, by the CNS. As ecdysone is synthesized and secreted by the ring gland, which lies outside of the CNS, and is converted to active 20E by peripheral organs (Petryk et al., 2003), (Huang et al., 2008), BBB-localized Mdr65 is in a prime position to regulate CNS-related behavioral sequences required for ecdysis and eclosion. When we investigated the cumulative timing of these behavioral transitions, we discovered that Mdr65 mutants took 24-hours longer to reach their final ecdysis/eclosion timepoint (Fig. 4A).

**Figure 4.**
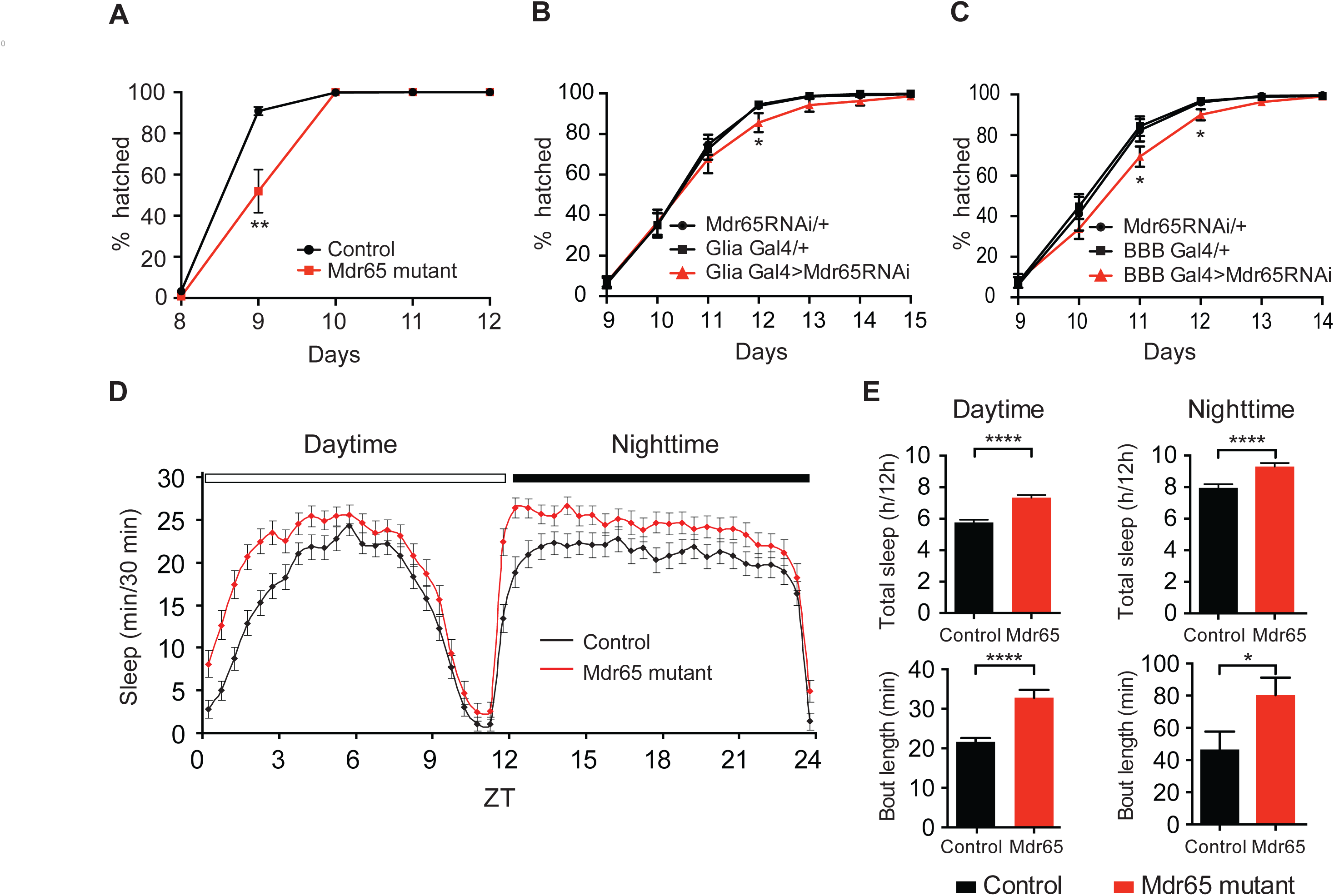
*Drosophila* xenobiotic efflux transporters can modulate behavior. (A) The cumulative percentage of adult flies hatching of wild type and Mdr65 null mutants. (B & C) The cumulative percentage of adult flies hatching following knock-down of Mdr65 in all glia (including the BBB) using the glial driver repo-Gal4 (B; Glia Gal4>Mdr65RNAi) or in the BBB using 9-137-Gal4 (C; BBB Gal4>Mdr65RNAi) compared to the heterozygote transgene controls. *, p<0.05. (D) The length of sleep bouts (min/30 minutes) for wild type and Mdr65 null mutants during the daytime and nighttime. (E) Total amount of sleep (hours/12 hour period) and average bout length during the daytime and nighttime. Error bars represent the SEM. T test *, p<0.05; ****, p≤0.0005.

To test whether BBB-localized Mdr65 can have cell autonomous behavioral roles, we assessed the effect of Mdr65 knock-down in the BBB. As the BBB is formed by glial cells in insects, we used pan-glial and BBB Gal4 drivers (DeSalvo et al., 2014) to express RNAi specific to Mdr65 in the BBB. Similar to the Mdr65 null mutants, but with a weaker effect, we found a statistically significant delay in adult eclosion (Fig. 4B&C). These data suggest that BBB-localized Mdr65 can regulate the behavioral transitions required for adult eclosion.

To determine whether xenobiotic efflux function can affect more complex behaviors in the adult animal, we assessed sleep behavior in Mdr65 mutants (Fig. 4D&E). It is well established that 20E has a dose-dependent effect on sleep/wake activity levels in *Drosophila;* increased amounts of 20E signaling in the mushroom body, a substructure that integrates and controls many behavioral functions of insects, is correlated with increased sleep intervals (Joiner et al., 2006, Pitman et al., 2006, Ishimoto and Kitamoto, 2010). We found that Mdr65 mutants demonstrate a significant increase in sleep behavior during both the daytime (27% increase) and nighttime (17% increase). Much of this increase is due to an increase in sleep bout length (Fig. 4E), a sleep characteristic that is influenced by 20E dosage (Ishimoto and Kitamoto, 2010). These data are consistent with increased 20E signaling in Mdr65 mutant brains, and suggests that BBB-localized Mdr65 can regulate complex behaviors.

### Mdr1 regulates anxiety-related behaviors in mice

Since Mdr65 loss of function alters behavior in flies, we tested whether loss of Mdr1 function in mice also has behavioral consequences. To do so, we tested adult male Mdr1 mutant mice against their littermate controls in a series of test that measure behaviors related to anxiety, motor control and social interaction.

In the novel environment exploration test Elevated Zero Maze (EZM), Mdr1 mutants spent less time in the open sections and more time in the closed sections of the EZM apparatus compared to controls (Fig. 5A). The behavior of Mdr1 mutants is consistent with an increase in anxiety levels, displaying caution when exploring a novel environment and preferring the more protected closed sections of the apparatus. Notably, the difference in behaviors between Mdr1 control and mutant mice is not due to abnormal movement or ambulation as Mdr1 controls and mutants scored similarly in these parameters (Fig. 5B).

**Figure 5.**
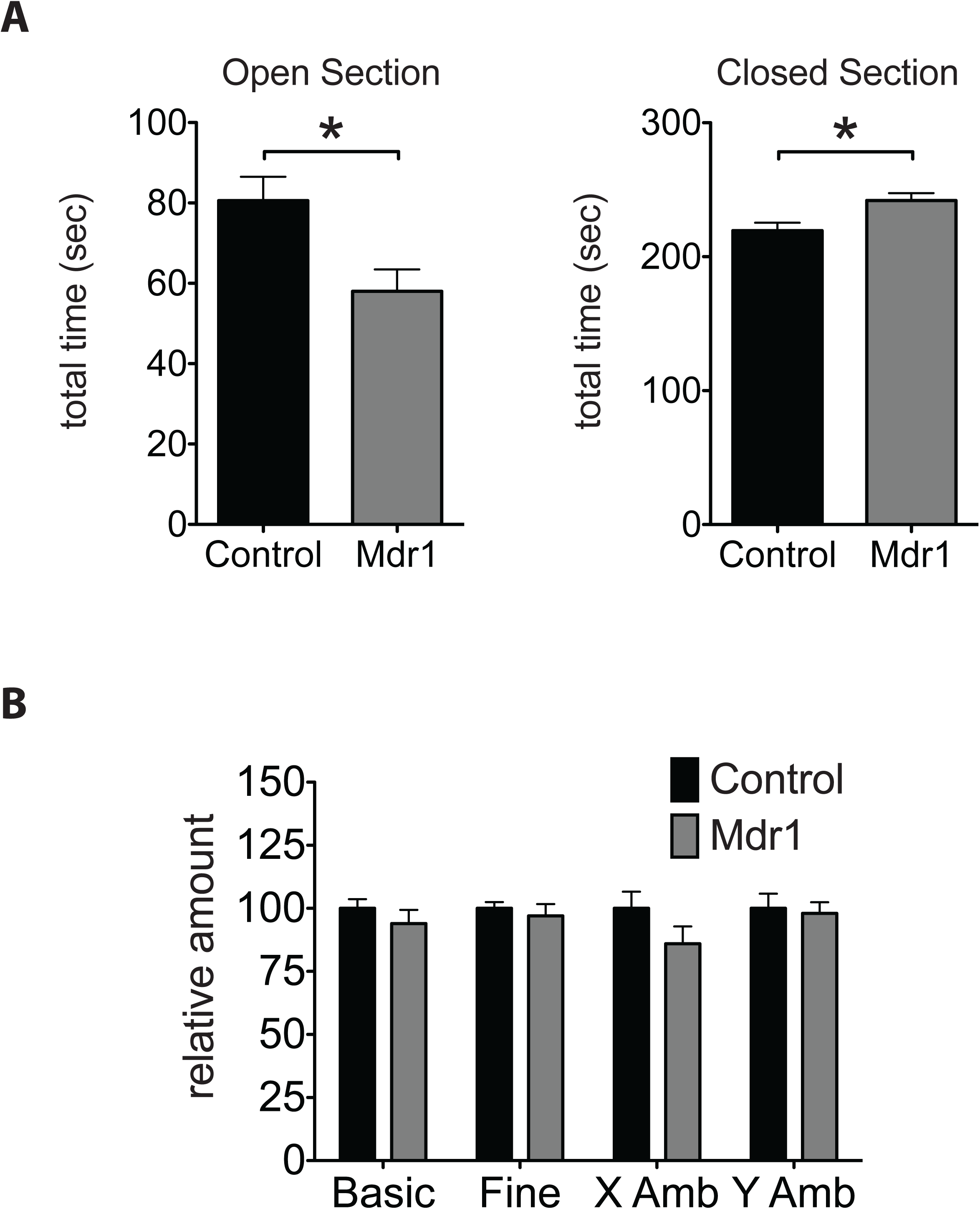
Anxiety-like behaviors are elevated in Mdr1 mutant mice. (A and B) Amount of time spent in the Open and Closed sections of the elevated zero maze (EZM) apparatus (A) and amount of movement and ambulation by adult littermate male Mdr1 control and mutant mice during the test (B). N= 16 controls and 9 Mdr1 mutant mice. Error bars represent SEM. Mann-Whitney * p<0.05.

Elevated anxiety in Mdr1 mutants was only observed when test mice (controls and mutants) were pre-exposed to the social interaction behavioral test, resident-intruder (RI) test (data not shown). Here, each test mouse was challenged in its home cage with a novel stimulus mouse. This gives the naïve test mice their first experience of a mouse that is not a parent or a sibling, leading to territorial, sexual and foraging competitive behaviors (Allen et al., 2010). The RI test also leads to displays of dominance and aggression, and experience of stress. These results suggest that once Mdr1 mutants experience competitiveness, dominance, aggression and/or stress from a novel male mouse, Mdr1 mutants experience above-normal anxiety in the EZM test. Similar to flies, these results show that BBB-enriched Mdr1 function can modulate complex animal behaviors. Moreover, these findings suggest that efflux transport by Mdr1 is important for inhibiting CNS access of anxiogenic molecules in adults or during the development of circuitry governing anxiety-related behaviors.

### An association between ABC efflux transporters and anxiety-related behaviors in humans

To expand on our findings that pharmacological inhibition of mouse Mdr1 can alter BBB partitioning of aldosterone, we asked whether pharmacological modulation of this transporter may influence behavior in humans.

Activity of human Mdr1 is subject to modulation by a variety of exogenous therapeutics and endogenous binding partners. Competitive binding interactions of drugs modulating Mdr1 may prevent it from interacting with its typical substrates, and lead to adverse effects. To investigate the possible role of human Mdr1 in regulating anxiety-related behaviors, we used an established analytical approach (Liu and Altman, 2015, Lounkine et al., 2012) to calculate Mdr1’s enrichment factor (EF) against each of 10,098 possible human adverse drug reactions (ADRs) gathered from the OFFSIDES collection of FDA drug reports (Tatonetti et al., 2012); Figure 6A).

**Figure 6.**
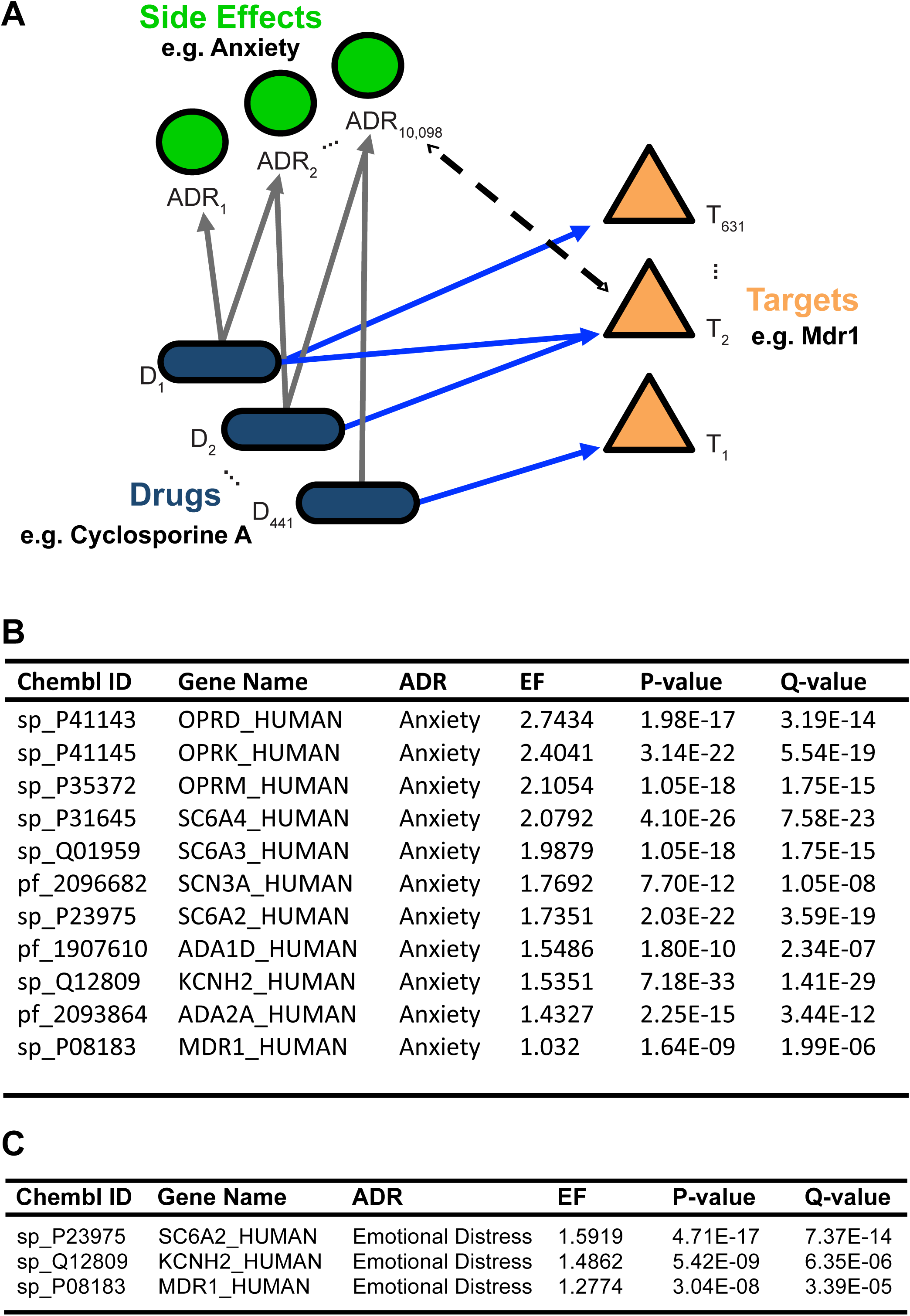
The adverse drug reaction Anxiety is linked with Mdr1-associated drugs. (A) Model diagram for target-ADR enrichment analysis. Enrichment factors were calculated from a unique set of 441 FDA-approved drugs, 10,098 adverse drug reactions (ADRs), and 631 CHEMBL targets after filtering (see Methods). Together, these analyses generated 1,018,208 EFs, of which only 2171 (0.2%) were statistically significant. (B & C) All target-ADR pairs significantly associated with the adverse drug reaction “Anxiety” or “Emotional Distress”. Targets are ordered by enrichment factor (EF) values, with a greater EF value being indicative of an increased number of observed target-ADR associations relative to the expected number of observations. Q-values indicate degree of confidence that the EF for each target-ADR association is not a false positive.

In our analysis, EFs quantify how strongly a therapeutic target (e.g. Mdr1) is associated with a particular ADR. To calculate EF values, we grouped drugs known to be inhibitors (n=13) or substrates (n=36) of Mdr1, as well as OFFSIDES compounds linked to Mdr1 via ChEMBL (Table 1), and examined whether these drugs, as a set, were enriched for certain ADRs e.g. “Anxiety” (UMLS code C00034E67) in comparison to drugs modulating other therapeutic targets. For a baseline of common drugs and their targets, we drew from 1104 FDA approved drugs and 1884 ChEMBL targets (Gaulton et al., 2012). After filtering (see Methods), this yielded 441 FDA-approved drugs organized into 631 ChEMBL targets, with an average of 3.97 drugs ± 0.21 per target. Out of the 1,018,208 unique target-ADR pairs represented by at least one drug within the dataset, a total of only 2171 pairs (0.2%) passed our statistical threshold (EF > 1.0, q-value < 1e-4; Table 2). Examining all targets linked to Anxiety, we found Mdr1 was one of only 12 targets significantly enriched (EF = 1.03, q-value = 1.80e-09, Fig. 6B). Furthermore, this enrichment was stable across multiple versions of ChEMBL (mean EF = 1.06 ± 0.02, Table 3).

**Table 1:**
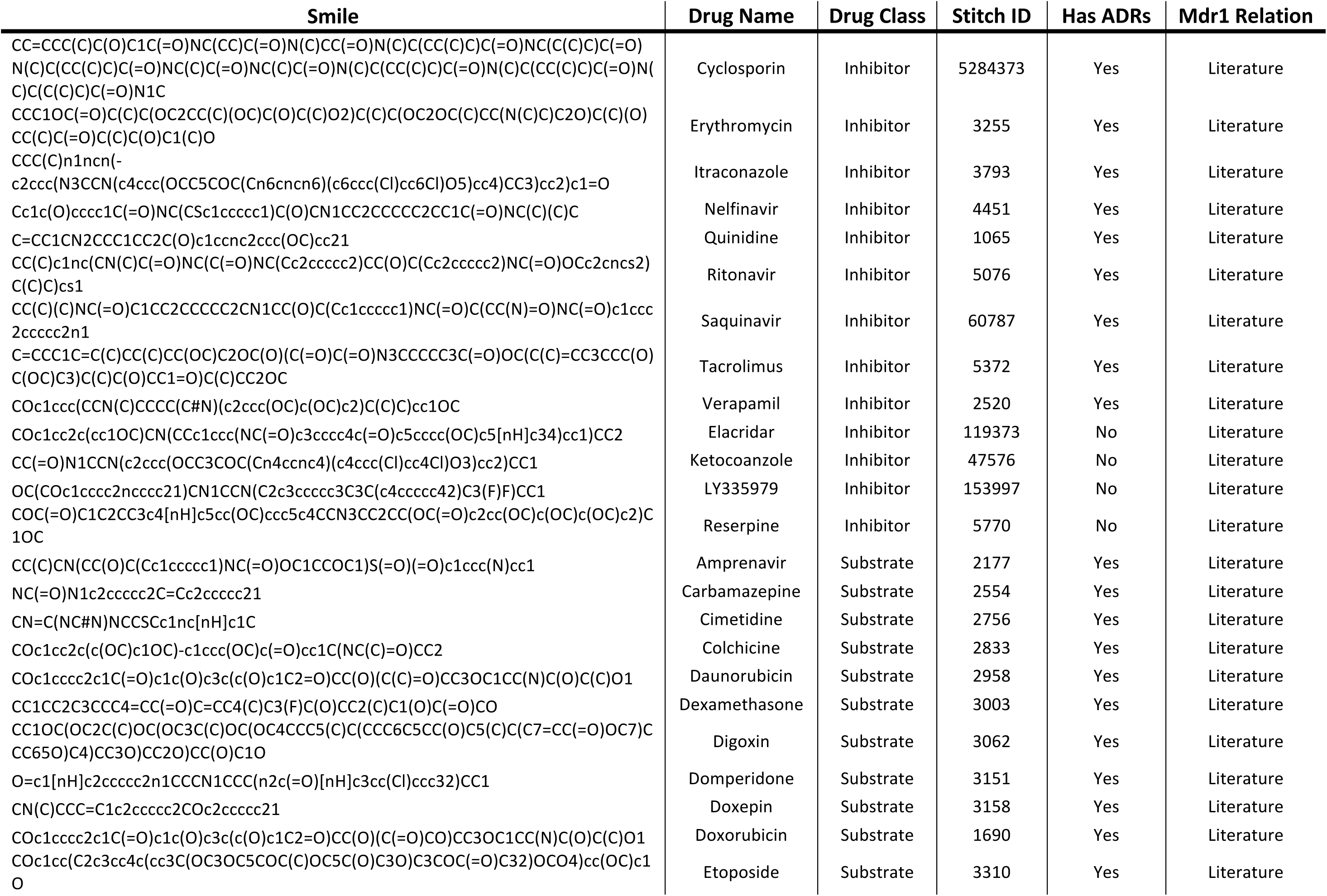
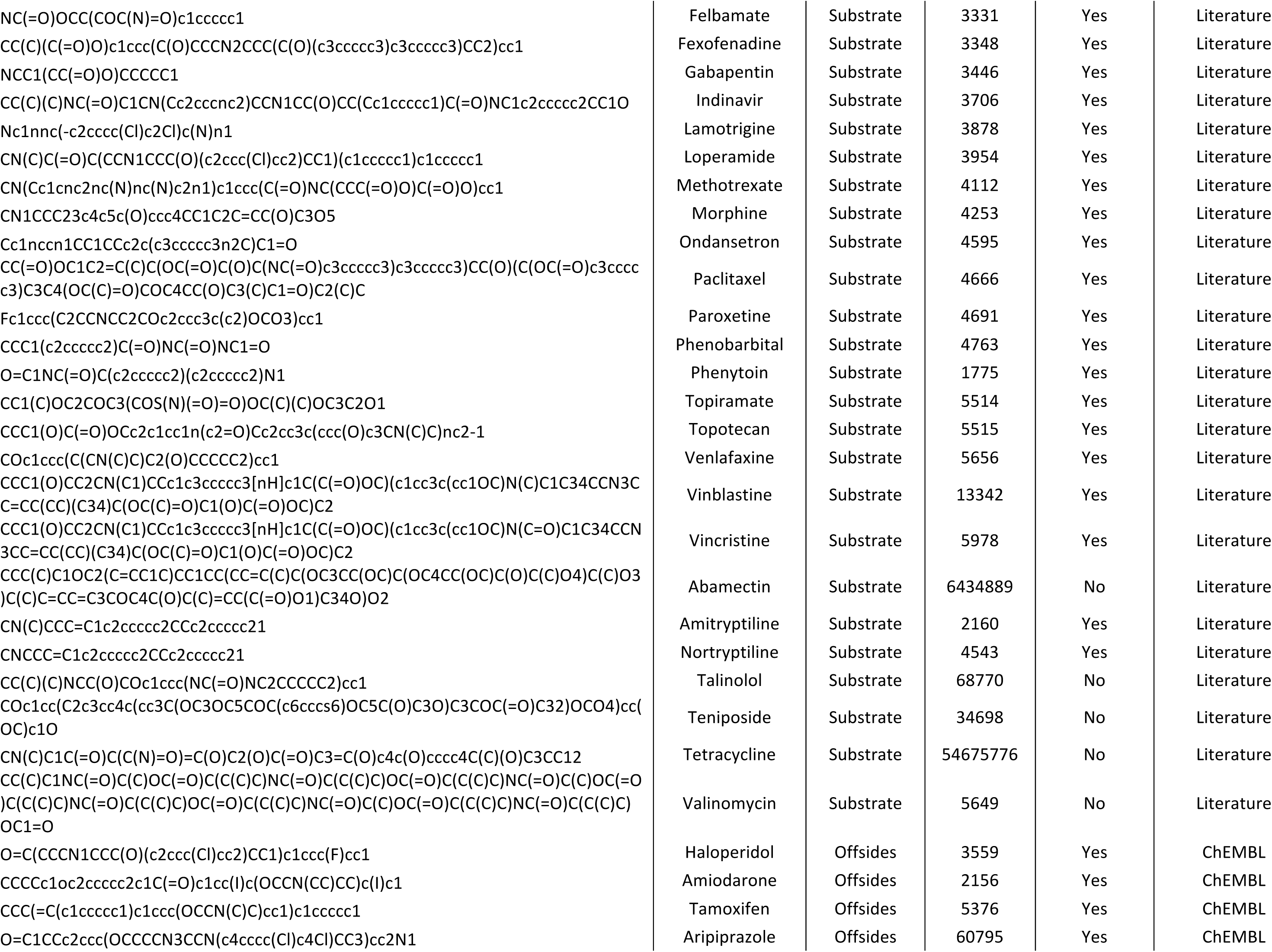
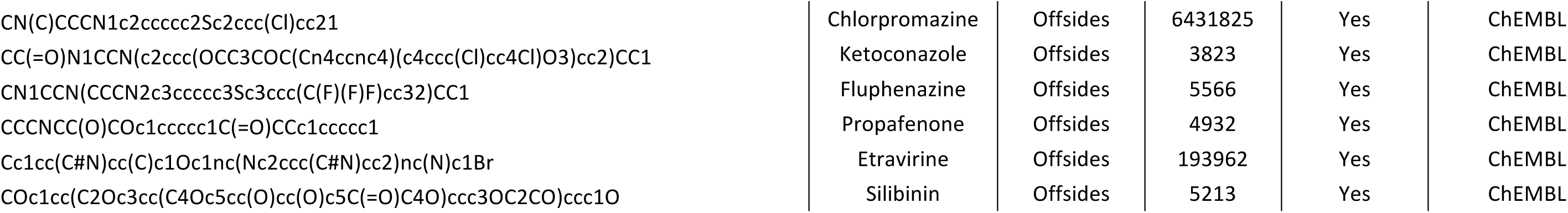
Inhibitors and Substrates of Mdr1.

**Table 2:** Top 100 Significantly Enriched Target-ADR Pairs.

| ChEMBL ID | Gene Name | Averse Drug Reaction | EF | P-value | Q-value |
| --- | --- | --- | --- | --- | --- |
| sp_P21917 | DRD4_HUMAN | neuroleptic malignant syndrome | 7.2812 | 1.28E-147 | 2.82E-144 |
| sp_P14416 | DRD2_HUMAN | neuroleptic malignant syndrome | 5.5261 | 2.49E-169 | 5.46E-166 |
| pf_2096682 | CHEMBL2096682 | pedal pulse absent | 5.1194 | 1.05E-22 | 1.83E-19 |
| sp_P14416 | DRD2_HUMAN | extrapyramidal disorder | 5.071 | 4.86E-158 | 1.07E-154 |
| sp_P35462 | DRD3_HUMAN | neuroleptic malignant syndrome | 5.0547 | 3.78E-142 | 8.30E-139 |
| sp_P14416 | DRD2_HUMAN | schizoffective disorder | 4.9192 | 3.17E-95 | 6.92E-92 |
| sp_P14416 | DRD2_HUMAN | Parkinson | 4.8826 | 1.82E-83 | 3.95E-80 |
| sp_P35462 | DRD3_HUMAN | increased appetite | 4.7492 | 1.17E-38 | 2.34E-35 |
| sp_P14416 | DRD2_HUMAN | Amphetamines | 4.6581 | 2.93E-109 | 6.40E-106 |
| sp_P35462 | DRD3_HUMAN | Parkinson | 4.5888 | 6.17E-62 | 1.31E-58 |
| sp_P08183 | MDR1_HUMAN | bone marrow depression | 4.587 | 7.73E-25 | 1.39E-21 |
| sp_P35462 | DRD3_HUMAN | amphetamines | 4.4839 | 5.62E-91 | 1.23E-87 |
| sp_P21917 | DRD4_HUMAN | agitated | 4.3326 | 1.34E-69 | 2.88E-66 |
| sp_P35462 | DRD3_HUMAN | extrapyramidal disorder | 4.3326 | 3.16E-97 | 6.92E-94 |
| sp_P23975 | SC6A2_HUMAN | nightmare | 4.3307 | 7.64E-49 | 1.58E-45 |
| sp_P14416 | DRD2_HUMAN | increased appetite | 4.2759 | 7.44E-45 | 1.53E-41 |
| sp_P35367 | HRH1_HUMAN | neuroleptic malignant syndrome | 4.2404 | 4.09E-129 | 8.97E-126 |
| sp_P08183 | MDR1_HUMAN | bacterial sepsis | 4.2047 | 4.78E-08 | 5.32E-05 |
| sp_P04278 | SHBG_HUMAN | adenocarcinoma | 4.1807 | 1.03E-39 | 2.07E-36 |
| sp_P35218 | CAH5A_HUMAN | blood potassium decreased | 4.0788 | 5.97E-44 | 1.22E-40 |
| sp_Q9Y2D0 | CAH5B_HUMAN | blood potassium decreased | 3.9846 | 4.34E-62 | 9.24E-59 |
| sp_P35462 | DRD3_HUMAN | delusion | 3.96 | 1.27E-85 | 2.76E-82 |
| sp_P08908 | 5HT1A_HUMAN | neuroleptic malignant syndrome | 3.9359 | 6.01E-67 | 1.29E-63 |
| sp_P08183 | MDR1_HUMAN | bone marrow disorder | 3.9005 | 4.97E-24 | 8.86E-21 |
| sp_P31645 | SC6A4_HUMAN | tardive dyskinesia | 3.8973 | 9.90E-121 | 2.17E-117 |
| sp_P14416 | DRD2_HUMAN | delusion | 3.8894 | 1.19E-91 | 2.60E-88 |
| pf_2094129 | CHEMBL2094129 | unresponsive to stimuli | 3.8572 | 1.10E-87 | 2.39E-84 |
| sp_P23975 | SC6A2_HUMAN | post traumatic stress disorder | 3.8547 | 1.26E-62 | 2.67E-59 |
| sp_P35367 | HRH1_HUMAN | agitated | 3.7848 | 2.41E-116 | 5.28E-113 |
| sp_P00915 | CAH1_HUMAN | blood potassium decreased | 3.7753 | 5.67E-28 | 1.06E-24 |
| sp_P35462 | DRD3_HUMAN | psychoses | 3.7137 | 4.69E-74 | 1.01E-70 |
| sp_P14416 | DRD2_HUMAN | tardive dyskinesia | 3.657 | 4.50E-80 | 9.74E-77 |
| sp_P31645 | SC6A4_HUMAN | disturbance in attention | 3.6455 | 3.47E-25 | 6.29E-22 |
| sp_P23280 | CAH6_HUMAN | blood potassium decreased | 3.6356 | 7.48E-25 | 1.35E-21 |
| sp_P35354 | PGH2_HUMAN | haematochezia | 3.6187 | 1.50E-35 | 2.96E-32 |
| pf_2093868 | CHEMBL2093868 | hyperhidrosis | 3.5833 | 1.41E-63 | 3.00E-60 |
| sp_P28223 | 5HT2A_HUMAN | psychoses | 3.5782 | 5.82E-59 | 1.23E-55 |
| sp_P35462 | DRD3_HUMAN | diabetic acidosis | 3.5557 | 7.55E-81 | 1.63E-77 |
| sp_P08908 | 5HT1A_HUMAN | carcinoid syndrome | 3.5423 | 9.21E-26 | 1.68E-22 |
| sp_P08183 | MDR1_HUMAN | lip erosion | 3.5068 | 7.04E-44 | 1.44E-40 |
| sp_P08183 | MDR1_HUMAN | mass excision | 3.5068 | 3.67E-26 | 6.73E-23 |
| sp_P35462 | DRD3_HUMAN | aura | 3.4722 | 1.14E-65 | 2.44E-62 |
| pf_2096682 | CHEMBL2096682 | aggression | 3.4671 | 1.69E-39 | 3.42E-36 |
| sp_P35367 | HRH1_HUMAN | diabetic acidosis | 3.4512 | 1.75E-85 | 3.80E-82 |
| sp_P31645 | SC6A4_HUMAN | carcinoid syndrome | 3.4297 | 8.96E-53 | 1.88E-49 |
| sp_P08183 | MDR1_HUMAN | alpha haemolytic streptococcal infection | 3.4231 | 1.10E-35 | 2.17E-32 |
| sp_Q01959 | SC6A3_HUMAN | disturbance in attention | 3.4018 | 2.19E-24 | 3.92E-21 |
| pf_2093864 | CHEMBL2093864 | Apnea | 3.3909 | 3.65E-59 | 7.73E-56 |
| sp_P31645 | SC6A4_HUMAN | drug addict | 3.3676 | 1.83E-54 | 3.85E-51 |
| pf_2093870 | CHEMBL2093870 | attempted suicide | 3.3509 | 5.86E-61 | 1.24E-57 |
| sp_P14416 | DRD2_HUMAN | psychoses | 3.3435 | 1.70E-65 | 3.63E-62 |
| pf_2093870 | CHEMBL2093870 | aura | 3.3097 | 1.83E-52 | 3.84E-49 |
| sp_Q96FL8 | S47A1_HUMAN | blood lactate dehydrogenase increased | 3.3091 | 4.86E-35 | 9.57E-32 |
| sp_P35367 | HRH1_HUMAN | blood triglycerides increased | 3.3028 | 2.23E-49 | 4.64E-46 |
| sp_P23975 | SC6A2_HUMAN | disturbance in attention | 3.2958 | 7.50E-54 | 1.57E-50 |
| sp_P35462 | DRD3_HUMAN | intention tremor | 3.2928 | 2.77E-21 | 4.76E-18 |
| sp_Q01959 | SC6A3_HUMAN | feeling abnormal | 3.2597 | 8.37E-25 | 1.51E-21 |
| pf_2093868 | CHEMBL2093868 | aura | 3.259 | 1.60E-54 | 3.36E-51 |
| sp_P35462 | DRD3_HUMAN | carcinoid syndrome | 3.2495 | 2.18E-52 | 4.55E-49 |
| pf_2095159 | CHEMBL2095159 | encephalopathy | 3.2425 | 7.63E-55 | 1.60E-51 |
| sp_P35354 | PGH2_HUMAN | gastric ulcer | 3.2416 | 6.17E-20 | 1.04E-16 |
| sp_P31645 | SC6A4_HUMAN | dysthymic disorder | 3.2371 | 2.74E-86 | 5.96E-83 |
| sp_P14416 | DRD2_HUMAN | aggression | 3.2352 | 5.76E-29 | 1.09E-25 |
| sp_P08908 | 5HT1A_HUMAN | diabetic acidosis | 3.2301 | 4.24E-39 | 8.52E-36 |
| sp_P14416 | DRD2_HUMAN | carcinoid syndrome | 3.2182 | 1.34E-49 | 2.78E-46 |
| sp_P14416 | DRD2_HUMAN | intention tremor | 3.2117 | 5.61E-24 | 9.97E-21 |
| sp_P31645 | SC6A4_HUMAN | feeling abnormal | 3.2021 | 1.19E-16 | 1.87E-13 |
| sp_P31645 | SC6A4_HUMAN | panic attack | 3.1943 | 1.90E-47 | 3.93E-44 |
| sp_P00918 | CAH2_HUMAN | azotaemia | 3.1863 | 8.25E-56 | 1.74E-52 |
| sp_P08183 | MDR1_HUMAN | gingival erosion | 3.1854 | 2.06E-32 | 3.99E-29 |
| sp_Q96FL8 | S47A1_HUMAN | neutrophil count decreased | 3.1776 | 2.19E-33 | 4.25E-30 |
| sp_Q01959 | SC6A3_HUMAN | carcinoid syndrome | 3.1739 | 1.31E-31 | 2.53E-28 |
| sp_P35462 | DRD3_HUMAN | agitated | 3.151 | 5.86E-84 | 1.27E-80 |
| sp_P35354 | PGH2_HUMAN | fibromyalgia | 3.1502 | 2.41E-39 | 4.85E-36 |
| sp_P23975 | SC6A2_HUMAN | carcinoid syndrome | 3.1409 | 9.70E-49 | 2.01E-45 |
| sp_P31645 | SC6A4_HUMAN | aggression | 3.1344 | 5.44E-58 | 1.15E-54 |
| sp_Q12809 | KCNH2_HUMAN | occult blood positive | 3.1272 | 5.64E-29 | 1.06E-25 |
| pf_2093864 | CHEMBL2093864 | renal tubular necrosis | 3.0925 | 8.43E-45 | 1.73E-41 |
| pf_1907610 | CHEMBL1907610 | Apnea | 3.091 | 3.56E-44 | 7.30E-41 |
| pf_2096682 | CHEMBL2096682 | blood glucose decreased | 3.0865 | 4.15E-33 | 8.07E-30 |
| sp_P08908 | 5HT1A_HUMAN | hyperhidrosis | 3.0707 | 1.00E-23 | 1.77E-20 |
| sp_P00918 | CAH2_HUMAN | blood potassium decreased | 3.0652 | 3.74E-23 | 6.57E-20 |
| sp_P08183 | MDR1_HUMAN | hepatic trauma | 3.0337 | 3.18E-17 | 5.07E-14 |
| sp_P08183 | MDR1_HUMAN | hodgkin's disease | 3.0337 | 1.48E-22 | 2.58E-19 |
| sp_P43166 | CAH7_HUMAN | angina | 3.025 | 1.74E-92 | 3.80E-89 |
| sp_P31645 | SC6A4_HUMAN | paranoia | 3.0147 | 1.07E-86 | 2.33E-83 |
| sp_P21917 | DRD4_HUMAN | drowsiness | 2.9822 | 2.28E-39 | 4.58E-36 |
| sp_P23975 | SC6A2_HUMAN | feeling abnormal | 2.9777 | 4.71E-39 | 9.44E-36 |
| sp_P14416 | DRD2_HUMAN | musculoskeletal stiffness | 2.9703 | 4.94E-46 | 1.02E-42 |
| sp_P28223 | 5HT2A_HUMAN | agitated | 2.9602 | 1.36E-42 | 2.77E-39 |
| sp_P23975 | SC6A2_HUMAN | major depression | 2.9427 | 7.35E-36 | 1.46E-32 |
| sp_P35367 | HRH1_HUMAN | intention tremor | 2.9297 | 2.17E-11 | 2.91E-08 |
| sp_P35367 | HRH1_HUMAN | Diabetic neuropathy | 2.9238 | 7.49E-36 | 1.48E-32 |
| pf_2095159 | CHEMBL2095159 | bradypnoea | 2.9145 | 4.59E-56 | 9.68E-53 |
| sp_P14416 | DRD2_HUMAN | acute brain syndrome | 2.9103 | 2.68E-42 | 5.45E-39 |
| sp_P14416 | DRD2_HUMAN | diabetic acidosis | 2.9103 | 1.09E-66 | 2.33E-63 |
| pf_1907610 | CHEMBL1907610 | blood pressure systolic increased | 2.8888 | 2.66E-42 | 5.41E-39 |
| sp_P28223 | 5HT2A_HUMAN | Diabetic neuropathy | 2.8785 | 6.86E-30 | 1.31E-26 |
| sp_P28335 | 5HT2C_HUMAN | diabetes mellitus non-insulin-dependent | 2.8782 | 1.32E-68 | 2.83E-65 |
| sp_P08183 | MDR1_HUMAN | radicular pain | 2.874 | 8.51E-17 | 1.34E-13 |

**Table 3:** Mdr1-Anxiety Enrichment Factors across different ChEMBL datasets.

| ChEMBL Version | Target ID | Gene Name | ADR | EF | P-value | Q-value |
| --- | --- | --- | --- | --- | --- | --- |
| 17 | sp_P08183 | MDR1_HUMAN | Anxiety | 1.0303 | 1.7953E-09 | 2.1777E-06 |
| 18 | CHEMBL4302 | MDR1_HUMAN | Anxiety | 1.1159 | 4.5643E-09 | 4.9317E-06 |
| 19 | CHEMBL4302 | MDR1_HUMAN | Anxiety | 1.0663 | 0.00074254 | 0.52646 |
| 20 | CHEMBL4302 | MDR1_HUMAN | Anxiety | 1.0693 | 0.0038258 | 1.0 |
| 21 | CHEMBL4302 | MDR1_HUMAN | Anxiety | 1.0592 | 2.9247E-09 | 5.0012E-06 |

While we were encouraged that patient-reported anxiety findings are associated with Mdr1, albeit with a subtle effect size, we acknowledge that anxiety-like feelings could be under-represented due to subjective behavioral reporting of drug side-effects. Likewise, the ADR term “Anxiety” may not thoroughly encompass the socially-primed effects observed in the mouse. Thus, we broadened the side-effect search terms to include “Emotional Distress” (UMLS code C0700361), which also classifies “Humiliation” (C0683285) and “Embarrassment” (C0679112). Strikingly, we found that Mdr1 was the 3^rd^ most enriched target for Emotional Distress (EF = 1.28, q value = 3.39e-05, Fig. 6C) out of all 1884 targets evaluated. Thus, human behavioral regulation by this Mdr1 pathway could be broad based, consistent with previous studies noting increased emotional stress in Mdr1 mutant mice (Schoenfelder et al., 2012).

To disprove the alternative hypothesis, we used the Similarity Ensemble Approach (SEA; (Keiser et al., 2007)) and confirmed that none of the 13 Mdr1 inhibitors had strongly predicted alternative off-targets more canonically associated with anxiety, such as the glucocorticoid or mineralocorticoid receptors. When expanding the analysis to include an additional 36 Mdr1 substrates, approximately 1.0% of the targets individually predicted by SEA could potentially be associated with Anxiety by literature search (Table 5). However, none of these alternative targets achieved significant enrichment with Anxiety by EF analysis (Fig. 6B).

These results, derived from human adverse drug reaction FDA data, lend support to our findings that modification of BBB xenobiotic efflux transporter function can alter the CNS concentrations of CNS-active endogenous molecules and disrupt behavioral pathways.

## Discussion

Using *Drosophila* and mouse studies, as well as bioinformatic analysis of human drug side effect data, we have firmly established a role for xenobiotic ABC transporters in modifying endobiotic localization. We show that disruption of BBB-enriched efflux transporter function leads to marked changes in CNS localization of endogenous steroid hormones and, more importantly, can influence neural functions that govern behavior. This is the first evidence of an evolutionarily conserved chemoprotective blood-brain-barrier function controlling an adult CNS physiology.

### The Blood-brain barrier controls partitioning of biologically potent endobiotics

This study highlights the role of the BBB in regulating CNS access of circulating steroid hormones. In particular, our fly data demonstrate that disruption of Mdr65 leads to an increase in the steroid 20E in the CNS. Extension of our analyses to mice establishes BBB-enriched Mdr1 as an important regulator of CNS aldosterone hormone levels. Together, these data suggest a critical role for the BBB in homeostatic maintenance of CNS steroid levels across species. As Mdr65 and Mdr1 have a broad spectrum of potential endogenous substrates (Fig. 1B), these findings implicate the BBB gateway in numerous small molecule regulatory events. This leads us to a hypothesis that altered endobiotic partitioning is sensed locally in the BBB, triggering compensatory mechanisms in an attempt to reinstate control (e.g. upregulation of Cyp18a1). This hypothesis is supported by the presence of the ecdysone receptor and aldosterone receptor transcripts in BBB cells (Daneman et al., 2010, DeSalvo et al., 2014), which may provide a signaling mechanism to translate changes in blood composition into BBB chemoprotective response and behavioral changes. In this model, the BBB participates actively as a brake or buffer on the overall CNS response to normal fluctuations of peripherally synthesized hormones and other CNS-active small molecules.

### ABC efflux transporters and the Blood-brain barrier are modulators of animal behaviors

Our *Drosophila* Mdr65 loss-of-function experiments revealed a significant role for ABC transporters in controlling CNS-related, ecdysone-regulated behaviors, including delayed developmental timing and increased sleep. Developmental progression is tightly regulated by precisely timed, ecdysone-induced behavioral cascades. Both the rise and fall in ecdysone levels are required for normal developmental transition. The rise in 20E primes the CNS by inducing the synthesis of ecdysis-related factors; however, the secretion of these factors to trigger the ecdysis behavioral cascade can only occur after the subsequent decline in 20E titer (Truman, 1981, Curtis et al., 1984, Truman, 1996, Zitnan et al., 1999, Zitnan et al., 2007); therefore, elevated CNS levels of ecdysone would be expected to cause a delay in this progression, like we see in Mdr65 mutants. Furthermore, feeding flies 20E causes an increased amount of sleep and, conversely, reducing ecdysone signaling in the brain causes a reduced amount of sleep (Ishimoto and Kitamoto, 2010). Our findings are consistent with an elevated CNS level of ecdysone causing increased sleep in Mdr65 mutant flies. A recent study showed a member of the ABCG class of transporters (atet) can control the vesicular release of ecdysone from the main steroidogenic organ (the ring gland) in the fly (Yamanaka et al., 2015). Depletion of atet from the ring gland led to a gross reduction in ecdysone release, leading to global loss of ecdysone availability. This resulted in a severe developmental arrest presumably due to the abolition of ecdysone pulses in the whole animal. Our results show that even subtle changes in BBB partitioning of steroid hormones can lead to significant changes in whole animal behavior, providing the BBB with a great capacity to modulate neural functions that govern processes as complex as sleep and anxiety.

*Drosophila* Mdr65 shares >40% sequence identity with mammalian Mdr1 and can transport Mdr1 substrates. In fact, human Mdr1 can rescue the chemoprotective BBB function of Mdr65 mutants, demonstrating that the xenobiotic-protective roles of BBB efflux transporters are conserved between flies and mammals (Mayer et al., 2009). Our data reveals that fly Mdr65 and mouse Mdr1 also have novel, conserved functions in regulating brain levels of endogenous steroids (20E in flies and aldosterone in mice) as well as steroid-related CNS-governed behaviors (developmental timing and sleep in flies, and anxiety-like behavior in mice). Moreover, we suggest that this novel function of ABC transporters may also manifest in human physiology.

### Pharmacological modification of ABC efflux transporters provides a potential mechanism for common adverse drug reactions

We showed that acute, pharmacological inhibition of Mdr1 can cause increased aldosterone in the brain. We suspected that unintended therapeutic modulation of Mdr1 activity may in turn misregulate endogenous substrate partitioning across the BBB, leading to unexpected side effects. Consistent with this, the unbiased analysis of 1104 FDA drugs, 1884 ChEMBL targets, and 10,098 ADRs revealed links to related behavioral effects of Mdr1 inhibition: that of anxiety and emotional distress.

Admittedly, the link between Mdr1 and Anxiety as a CNS side effect in humans is subtle (Fig. 6B). While in line with our observation that modification of ABC transporters has significant CNS behavioral effects in the fly and mouse, this initial human result merits further exploration as its enrichment factor reflected several possible limitations. First, Anxiety is a widely prevalent ADR appearing for a diverse set of drugs (478 drugs: (43% of all drugs from OFFSIDES dataset). Second, few of the 1104 drugs in the dataset have been definitively characterized with respect to Mdr1, potentially confounding the analysis as unreported substrate or inhibitor roles would be expected to weaken the calculated enrichment of Mdr1 to Anxiety. Third, Mdr1 itself exhibits a broad substrate profile, as evidenced by the large number of disparate ADRs to which it is linked (216; Table 4). Although we note that only 6% of its associated ADRs might be considered behaviorally relevant, all are taken into consideration when calculating enrichment, which can weaken Mdr1-related scores. Nevertheless, we were intrigued to see the Mdr1-Anxiety link emerge from this pan-ADR analysis at all, given these limitations. The strong enrichment between Mdr1 and Emotional Distress bypasses many of these limitations, and lends considerable support to Mdr1’s potential role in stress-related behaviors, particularly those with a social aspect such as Humiliation. This may be consistent with the resident-intruder test priming effect observed in Mdr1 mutant mice, but further study would be required to investigate the potential roles of dominance and aggression in this context.

**Table 4:** Top 100 Adverse Drug Events Significantly Enriched for ChEMBL Target Mdr1.

| ChEMBL ID | Uniprot_ID | Adverse Drug Reaction | EF-value | P-value | Q-value |
| --- | --- | --- | --- | --- | --- |
| sp_P08183 | MDR1_HUMAN | bone marrow depression | 4.5945 | 7.02E-25 | 1.28E-21 |
| sp_P08183 | MDR1_HUMAN | bacterial sepsis | 4.2542 | 3.27E-08 | 3.68E-05 |
| sp_P08183 | MDR1_HUMAN | bone marrow disorder | 3.8288 | 7.02E-25 | 1.28E-21 |
| sp_P08183 | MDR1_HUMAN | lip erosion | 3.5126 | 6.02E-44 | 1.25E-40 |
| sp_P08183 | MDR1_HUMAN | mass excision | 3.5126 | 9.11E-28 | 1.72E-24 |
| sp_P08183 | MDR1_HUMAN | alpha haemolytic streptococcal infection | 3.4545 | 8.07E-38 | 1.63E-34 |
| sp_P08183 | MDR1_HUMAN | gingival erosion | 3.1906 | 6.15E-34 | 1.22E-30 |
| sp_P08183 | MDR1_HUMAN | hodgkin's disease | 3.0549 | 8.24E-23 | 1.46E-19 |
| sp_P08183 | MDR1_HUMAN | hepatic trauma | 3.0387 | 2.95E-17 | 4.72E-14 |
| sp_P08183 | MDR1_HUMAN | radicular pain | 2.8788 | 5.92E-18 | 9.61E-15 |
| sp_P08183 | MDR1_HUMAN | brain natriuretic peptide increased | 2.8121 | 1.02E-20 | 1.76E-17 |
| sp_P08183 | MDR1_HUMAN | non-hodgkin's lymphoma | 2.7527 | 5.08E-09 | 5.96E-06 |
| sp_P08183 | MDR1_HUMAN | electrocardiogram qrs complex prolonged | 2.7348 | 3.57E-26 | 6.61E-23 |
| sp_P08183 | MDR1_HUMAN | hyperpigmentation | 2.7348 | 1.77E-13 | 2.56E-10 |
| sp_P08183 | MDR1_HUMAN | hyperparathyroidism | 2.6775 | 7.47E-16 | 1.16E-12 |
| sp_P08183 | MDR1_HUMAN | subcutaneous nodule | 2.6224 | 2.77E-15 | 4.22E-12 |
| sp_P08183 | MDR1_HUMAN | lymphoma | 2.6021 | 7.96E-27 | 1.49E-23 |
| sp_P08183 | MDR1_HUMAN | polyp | 2.5924 | 4.77E-49 | 1.00E-45 |
| sp_P08183 | MDR1_HUMAN | joint contracture | 2.5812 | 1.99E-18 | 3.28E-15 |
| sp_P08183 | MDR1_HUMAN | subclavian vein thrombosis | 2.5696 | 7.47E-35 | 1.49E-31 |
| sp_P08183 | MDR1_HUMAN | crackles lung | 2.5525 | 1.78E-13 | 2.58E-10 |
| sp_P08183 | MDR1_HUMAN | nephrogenic fibrosing dermopathy | 2.53 | 4.26E-29 | 8.14E-26 |
| sp_P08183 | MDR1_HUMAN | blood parathyroid hormone increased | 2.4862 | 3.55E-26 | 6.57E-23 |
| sp_P08183 | MDR1_HUMAN | hypertonic bladder | 2.3781 | 6.06E-22 | 1.06E-18 |
| sp_P08183 | MDR1_HUMAN | local swelling | 2.2633 | 8.82E-38 | 1.78E-34 |
| sp_P08183 | MDR1_HUMAN | synovitis | 2.2522 | 1.47E-24 | 2.66E-21 |
| sp_P08183 | MDR1_HUMAN | bone scan abnormal | 2.239 | 1.05E-16 | 1.65E-13 |
| sp_P08183 | MDR1_HUMAN | enlarged spleen | 2.2294 | 6.06E-20 | 1.03E-16 |
| sp_P08183 | MDR1_HUMAN | heart valve incompetence | 2.2059 | 2.00E-16 | 3.13E-13 |
| sp_P08183 | MDR1_HUMAN | ear infection | 2.205 | 9.32E-10 | 1.15E-06 |
| sp_P08183 | MDR1_HUMAN | elevated triglycerides | 2.1879 | 1.32E-19 | 2.24E-16 |
| sp_P08183 | MDR1_HUMAN | fib | 2.0585 | 1.37E-22 | 2.42E-19 |
| sp_P08183 | MDR1_HUMAN | skin plaque | 2.0475 | 3.85E-14 | 5.69E-11 |
| sp_P08183 | MDR1_HUMAN | Adenopathy | 2.0233 | 4.61E-20 | 7.84E-17 |
| sp_P08183 | MDR1_HUMAN | carotid artery stenosis | 2.0151 | 2.29E-11 | 3.07E-08 |
| sp_P08183 | MDR1_HUMAN | diastolic dysfunction | 2.0151 | 2.55E-11 | 3.41E-08 |
| sp_P08183 | MDR1_HUMAN | Hepatitis C | 2.0151 | 2.66E-19 | 4.49E-16 |
| sp_P08183 | MDR1_HUMAN | Scar | 2.0034 | 2.62E-29 | 5.01E-26 |
| sp_P08183 | MDR1_HUMAN | cirrhosis | 1.996 | 1.74E-19 | 2.94E-16 |
| sp_P08183 | MDR1_HUMAN | abscess drainage | 1.9804 | 1.99E-18 | 3.28E-15 |
| sp_P08183 | MDR1_HUMAN | fatty liver | 1.9745 | 2.73E-18 | 4.48E-15 |
| sp_P08183 | MDR1_HUMAN | Rhinitis | 1.9724 | 3.01E-18 | 4.93E-15 |
| sp_P08183 | MDR1_HUMAN | joint stiffness | 1.9596 | 4.86E-20 | 8.25E-17 |
| sp_P08183 | MDR1_HUMAN | osteoporosis | 1.9503 | 6.39E-18 | 1.04E-14 |
| sp_P08183 | MDR1_HUMAN | cervicobrachial syndrome | 1.9468 | 6.55E-15 | 9.91E-12 |
| sp_P08183 | MDR1_HUMAN | back disorder | 1.924 | 2.73E-10 | 3.49E-07 |
| sp_P08183 | MDR1_HUMAN | pseudomonas infection | 1.92 | 2.40E-36 | 4.83E-33 |
| sp_P08183 | MDR1_HUMAN | haematochezia | 1.9197 | 7.63E-16 | 1.18E-12 |
| sp_P08183 | MDR1_HUMAN | pancreatitis chronic | 1.8976 | 1.58E-34 | 3.14E-31 |
| sp_P08183 | MDR1_HUMAN | skin induration | 1.883 | 2.33E-17 | 3.75E-14 |
| sp_P08183 | MDR1_HUMAN | gingival oedema | 1.8677 | 2.42E-10 | 3.11E-07 |
| sp_P08183 | MDR1_HUMAN | ear ache | 1.8631 | 6.79E-11 | 8.96E-08 |
| sp_P08183 | MDR1_HUMAN | Tendinitis | 1.8503 | 2.74E-16 | 4.28E-13 |
| sp_P08183 | MDR1_HUMAN | blood amylase increased | 1.8467 | 4.47E-10 | 5.62E-07 |
| sp_P08183 | MDR1_HUMAN | spinning sensation | 1.8467 | 1.71E-12 | 2.38E-09 |
| sp_P08183 | MDR1_HUMAN | Acute Respiratory Distress Syndrome | 1.8432 | 4.70E-55 | 1.00E-51 |
| sp_P08183 | MDR1_HUMAN | febrile neutropenia | 1.8232 | 7.75E-10 | 9.60E-07 |
| sp_P08183 | MDR1_HUMAN | blood sodium increased | 1.8089 | 9.04E-40 | 1.84E-36 |
| sp_P08183 | MDR1_HUMAN | Intervertebral Disc Herniation | 1.7771 | 1.02E-19 | 1.73E-16 |
| sp_P08183 | MDR1_HUMAN | ischias | 1.7713 | 3.84E-14 | 5.67E-11 |
| sp_P08183 | MDR1_HUMAN | abdominal distension | 1.7403 | 5.28E-11 | 7.01E-08 |
| sp_P08183 | MDR1_HUMAN | cervical spinal stenosis | 1.7332 | 2.88E-09 | 3.44E-06 |
| sp_P08183 | MDR1_HUMAN | angiitis | 1.7325 | 3.86E-12 | 5.33E-09 |
| sp_P08183 | MDR1_HUMAN | dental pain | 1.7169 | 9.45E-32 | 1.85E-28 |
| sp_P08183 | MDR1_HUMAN | hydronephrosis | 1.7042 | 4.67E-12 | 6.42E-09 |
| sp_P08183 | MDR1_HUMAN | basal cell carcinoma | 1.6954 | 3.71E-08 | 4.15E-05 |
| sp_P08183 | MDR1_HUMAN | cyst | 1.6923 | 3.99E-19 | 6.68E-16 |
| sp_P08183 | MDR1_HUMAN | lumbar spinal stenosis | 1.6892 | 8.36E-12 | 1.14E-08 |
| sp_P08183 | MDR1_HUMAN | spinal disorder | 1.6867 | 7.45E-09 | 8.68E-06 |
| sp_P08183 | MDR1_HUMAN | Sinusitis | 1.6852 | 3.30E-21 | 5.72E-18 |
| sp_P08183 | MDR1_HUMAN | Scoliosis | 1.6842 | 6.78E-35 | 1.35E-31 |
| sp_P08183 | MDR1_HUMAN | stem cell transplant | 1.6744 | 2.06E-10 | 2.66E-07 |
| sp_P08183 | MDR1_HUMAN | rash generalised | 1.6707 | 4.10E-10 | 5.16E-07 |
| sp_P08183 | MDR1_HUMAN | bursitis | 1.6604 | 7.00E-13 | 9.95E-10 |
| sp_P08183 | MDR1_HUMAN | Candida Infection | 1.6527 | 1.53E-11 | 2.07E-08 |
| sp_P08183 | MDR1_HUMAN | thyroid neoplasia | 1.6503 | 1.26E-14 | 1.88E-11 |
| sp_P08183 | MDR1_HUMAN | burning sensation | 1.6432 | 9.91E-19 | 1.65E-15 |
| sp_P08183 | MDR1_HUMAN | joint range of motion decreased | 1.6373 | 1.68E-12 | 2.34E-09 |
| sp_P08183 | MDR1_HUMAN | birth defect | 1.6272 | 5.02E-14 | 7.37E-11 |
| sp_P08183 | MDR1_HUMAN | bone inflammation | 1.6243 | 1.11E-08 | 1.27E-05 |
| sp_P08183 | MDR1_HUMAN | Incontinence | 1.6188 | 1.83E-10 | 2.38E-07 |
| sp_P08183 | MDR1_HUMAN | musculoskeletal stiffness | 1.6137 | 4.93E-54 | 1.05E-50 |
| sp_P08183 | MDR1_HUMAN | oropharyngeal pain | 1.6121 | 2.07E-08 | 2.35E-05 |
| sp_P08183 | MDR1_HUMAN | leukoencephalopathy | 1.6108 | 1.14E-09 | 1.40E-06 |
| sp_P08183 | MDR1_HUMAN | reflux esophagitis | 1.6108 | 1.14E-09 | 1.40E-06 |
| sp_P08183 | MDR1_HUMAN | asthmatic crisis | 1.6087 | 1.17E-14 | 1.76E-11 |
| sp_P08183 | MDR1_HUMAN | bone debridement | 1.6071 | 8.75E-11 | 1.15E-07 |
| sp_P08183 | MDR1_HUMAN | rib fracture | 1.5875 | 1.74E-08 | 1.98E-05 |
| sp_P08183 | MDR1_HUMAN | haemodialysis | 1.5865 | 4.20E-10 | 5.29E-07 |
| sp_P08183 | MDR1_HUMAN | acute leukaemia | 1.5821 | 5.70E-23 | 1.01E-19 |
| sp_P08183 | MDR1_HUMAN | skin tightness | 1.5708 | 8.89E-11 | 1.17E-07 |
| sp_P08183 | MDR1_HUMAN | gall bladder | 1.5692 | 9.27E-12 | 1.26E-08 |
| sp_P08183 | MDR1_HUMAN | Caesarean Section | 1.5628 | 8.37E-13 | 1.18E-09 |
| sp_P08183 | MDR1_HUMAN | skin lesion | 1.5497 | 6.41E-08 | 7.02E-05 |
| sp_P08183 | MDR1_HUMAN | hyperbaric oxygen therapy | 1.5193 | 5.87E-08 | 6.46E-05 |
| sp_P08183 | MDR1_HUMAN | streptococcal infection | 1.5193 | 2.59E-10 | 3.32E-07 |
| sp_P08183 | MDR1_HUMAN | hypoaesthesia | 1.5175 | 6.57E-25 | 1.20E-21 |
| sp_P08183 | MDR1_HUMAN | primary sequestrum | 1.5099 | 5.00E-09 | 5.88E-06 |
| sp_P08183 | MDR1_HUMAN | gastric hyperplasia | 1.4999 | 8.84E-14 | 1.29E-10 |
| sp_P08183 | MDR1_HUMAN | fistula | 1.4982 | 5.97E-14 | 8.76E-11 |

These Drug-Target-ADR enrichment factor results are supported by the confluence of complementary computational and *in vivo* lines of evidence we have presented here. The role of drugs in modulating the CNS entry of endogenous steroids, as shown by our co-injections of CsA and aldosterone, and their effects on behavior suggest a new way of thinking about complex drug-drug interactions (Zhang and Sparreboom, 2017).

Of note, pharmacological inhibition of Mdr1 yields no more than a 2-fold change in drug partitioning at the BBB; therefore, CNS adverse effects might seem unlikely in this context (Kalvass et al., 2013). However, our data showing that pharmacological inhibition of Mdr1 in vivo can lead to a 3.3-fold increase in brain aldosterone levels suggest that effects on endogenous molecule partitioning may be greater than previously predicted, at least transiently. Indeed, the most prevalent adverse drug reactions– altered sleep/wake cycles, depression and anxiety– run the gamut of drug classes and chemical subtypes, with no obvious common mechanism. It is possible that the same mechanisms that evolved to regulate CNS access of endogenous molecules are repurposed by the BBB to chemically isolate xenobiotics from the brain; hence, a tension occurs between endobiotic and xenobiotic partitioning during drug exposure. ABC efflux transporters are common drug targets; thus, unintended transporter-mediated modulation of endobiotic partitioning merits further investigation as a common mechanism for such wide-spread adverse drug reactions.

In addition, genetic and environmental conditions altering human ABC transporter function may pose a previously unappreciated risk to human health. Human Mdr1 function is altered in many CNS pathologies, and single nucleotide polymorphisms (SNPs) can affect its function (Jeong et al., 2007),(Pauli-Magnus and Kroetz, 2004). For example, expression and activity levels of ABC transporters, including Mdr1, are altered in certain neurodegenerative disorders, such as Alzheimer’s disease, Parkinson’s disease and Creutzfeld-Jacob disease, and in normal aging (Vogelgesang et al., 2002), (Loscher and Potschka, 2005b), (Vogelgesang et al., 2006, Bauer et al., 2009, Vautier and Fernandez, 2009, Jablonski et al., 2014). Furthermore, behavioral symptoms, like anxiety and depression, are commonly associated with these disorders (Chemerinski et al., 1998). Thus, altered endobiotic pharmacokinetics at the BBB may provide a novel mechanism for these clinical manifestations.

Our study shows that BBB-enriched chemoprotective transporters are key regulators of CNS levels of biologically potent endogenous molecules, and animal behavior. In addition we show that pharmacological inhibition of Mdr1 also leads to an increase in CNS steroid levels, and is associated with the ADR anxiety and emotional distress in humans. Thus, we suggest that increased partition of endobiotics into the brain affecting anxiety-related behaviors is a novel and logical outcome of the pharmacologic inhibition of normal BBB chemoprotective function by drugs at steady state. Furthermore, we propose that the BBB can be considered as a target tissue for regulating behavior and not just a conduit and obstacle for small molecule partition into brain space. Further study of BBB regulated processes may yield other roles for BBB function in behavior, and new systems for targeting behavioral therapy that are more accessible than internal brain pathways.

## Materials and Methods

### Homology Modeling

The protein sequence of Mdr65 was obtained from the Uniprot database, accession number Q000748. The structure of mouse Mdr1 with docked saquinavir sharing 40% sequence identity with Mdr65, was used as the template (PDB ID 3G60). Sequence alignment was performed using BLAST, and the Mdr65 model was built using the standard homology protocol within the Prime module of Schrodinger Suite 2012. The model was subsequently processed in the Prepwizard module to optimize hydrogens; the model was subsequently subjected to restrained energy minimization with the saquinavir ligand removed.

### Ligand Docking

Flexible receptor docking was performed using the induced fit docking protocol developed previously (Dolghih et al., 2011) and final scoring was implemented using the extra precision (XP) Glide with the OPLS2005 force field. A 10 × 10 × 10 Å inner docking box with the center at coordinates (19.0, 46.0, -6.0) Å was used. The number of poses saved during the initial docking was set to 100. In the next stage, only residues within 5 Å of each ligand were minimized and up to 20 top poses were saved and scored with Glide XP.

### Ligands

Ligand structures were obtained from the KEGG database of biologically relevant molecules and processed using the *Ligprep* 2.4 module. Parameters were assigned based on the OPLS2005 force field. Ionization states were assigned by *Epik*, and groups with pKa between 5 and 9 were treated as neutral while those outside the range were treated as charged.

### *Drosophila* genetics

The following fly strains were used during this study: Canton-S and isogenized w^-1118^ (w-ISO) wild type strains, the EcRLBD ecdysone reporter stock #23656, UAS-StingerGFP (generated from stock #28863), and the glial cell driver repo-Gal4 stock #7415 (from the Bloomington Drosophila Stock Center, Indiana University, Bloomington, IN), the pMdr65 null mutant strain (KG08723 from the Drosophila Genetic Resource Center, Kyoto Institute of Technology, Kyoto, Japan), the UAS-Mdr65RNAi Stock #9019 (from the Vienna Drosophila Research Center (Dietzl et al., 2007), and 9-137 Gal4 (DeSalvo et al., 2014).

### *In vivo* competitive efflux transport assay

Male flies (4-8 days old) were hemolymph-injected (Mayer et al., 2009) with 200 ng Rhodamine B (R6626; Sigma) and either 20% ethanol (vehicle) or approximately 500 ng (∼100 nl of 4.8 mg/ml) 20-hydroxyecdysone (20-E; H5142; Sigma). Flies were left for 2 h followed by rapid dissection of the brains in PBS (<2 min per dissection). The brains were quickly rinsed in PBS and immediately placed in individual wells of a foil-covered fluorometer plate containing 200 μl 10% SDS. Brains were left to dissociate for 30 min followed by fluorescence reading on a fluorometer (ex. 535 nm/ em. 595 nm). Brain fluorescence readings were normalized to whole body readings to account for individual differences in dye loading and elimination from the fly. N=7 (except for n=4 for the Mdr65 RhoB/20E condition). Graphpad Prism software was used to perform ANOVA statistical analyses after removal of one statistically identified outlier in the wild type RhoB/Vehicle condition (Dixon test ratio =0.726, critical value =0.507, no other outliers were identified).

### Western blotting

For the EcRLBD reporter time-course assays, male flies (4-8 days old) were hemolymph-injected (Mayer et al., 2009) with 20% ethanol (vehicle) or 4.8 mg/ml 20-E and were frozen after specified time-points. Heads were separated from the bodies and the heads were homogenized in sample buffer (0.04% Bromophenol Blue, 4% SDS, 4% glycerol, 10% β-mercaptoethanol, 0.1M Tris, pH 6.8). The equivalent of two fly heads was run for each time-point. Western analysis was performed using standard 10% PAGE gels blotted onto PVDF membranes (BioRad). Membranes were blocked using 5% non-fat milk in Tris-buffered saline supplemented with 0.1% Tween-20 (0.1% TBS-T) for 1 h. Primary antibody hybridization occurred overnight at 4°C followed by 3x washes in 0.1% TBS-T, secondary antibody hybridization for 1 h, then 3x washes in 0.1% TBS-T. Primary antibodies used included rabbit α-GFP (1:1000; ab6556; Abcam) and mouse α-tubulin (1:100; E7; DSHB). Secondary antibodies used included goat α-rabbit HRP (1:40,000; #28177; AnaSpec) and goat α-mouse HRP (1:1000; #31430; Pierce).

### Immunofluorescence

Male flies (4-8 days old) were hemolymph-injected (Mayer et al., 2009) with 12.5 μg/μl 70 kDa Texas Red dextran (D1864; Invitrogen) and either 20% ethanol (vehicle) or approximately 500 ng 20-hydroxyecdysone and left to recover overnight (approximately 16 h). Whole heads were bisected from the bodies, the proboscis removed, and the heads fixed in 3.7% paraformaldehyde/PBS for 15 min. Brains were dissected in PBS, removing all the fat bodies and large trachea but taking care to not damage the BBB. Brains were washed in PBS followed by incubation in blocking buffer (5% normal calf serum (NCS) or donkey serum, 4% Tween-20 in PBS) for 1 h, then primary antibody incubation overnight at 4°C. Brains were washed in PBS (3x 30 min) followed by incubation in secondary antibody for 45 min. Brains were washed in PBS (3x 45 min) and mounted in Dako fluorescent mounting medium (S3023; Dako) on glass slides with nail polish posts. Brains were imaged immediately. Primary antibodies used include rabbit α-GFP polyclonal (1:1000; ab6556; Abcam), mouse α-repo monoclonal (1:10; 8D12; Developmental Studies Hybridoma Bank (DSHB)). Secondary antibodies used include goat α-rabbit FITC (1:100; Invitrogen), goat α-mouse Cy5 (1:1000; ab6563; Abcam), donkey α-rabbit Alexa Fluor 488 (1:1000; Invitrogen), donkey α-mouse Alexa Fluor 647 (1:1000; Invitrogen). Images were captured of the optic lobe region of the CNS using a Zeiss LSM 510 confocal microscope. Confocal z-section images were acquired every 0.5 μm from the BBB surface through to the cross-section. Confocal settings were maintained across all samples within an experiment. At least 3 brains were imaged per condition, and data analyzed from at least 3 technical replicates.

### Image analyses

Quantification of the number of GFP-positive cells in the CNS of Mdr65 mutants and controls was calculated using ImageJ. Z-stack sections were imported as TIFF files, converted to binary (8-bit) and identical threshold settings were applied to all z-stacks. Data were normalized to the number of GFP-positive cells at the brain surface.

### Developmental timing and eclosion

Vials containing 20 males and females were transferred onto fresh food with added yeast paste every 2 h at 25°C to allow egg laying. Flies were maintained at 25°C except when scoring. The number of hatched flies was scored daily. At least 2 technical replicates were performed. For the BBB knock-down of Mdr65 (using repo Gal4 and 9-137 Gal4), the eclosion assays were performed in bottles and the flies were left to lay eggs for approximately 2 days. The number of hatched flies was scored twice daily and the numbers pooled per day. At least 2 technical replicates were performed.

### Sleep analysis

Newly eclosed flies were collected under brief CO_2_ anesthesia over a three-day period. Males were housed with 10 flies per vial on standard cornmeal agar media, and aged for 3-5 days prior to experimentation. For sleep analysis, flies were individually aspirated into a glass tube (5 [W] × 65 [L] mm) with regular fly food, and their locomotor activity was monitored using the Drosophila Activity Monitoring (DAM) system (Trikinetics, Waltham, MA, USA) while subjected to 12 hr light and 12 hr dark cycles at 25°C with 65% humidity. Flies were acclimated to the experimental conditions for one day before sleep was analyzed. Locomotor activity data were collected at 1-min intervals for 3 days, and analyzed with a Microsoft (Redmond, WA, USA) Excel-based macro program as described previously (Hendricks et al., 2003),(Kume et al., 2005). A sleep bout was defined as 5 or more minutes of behavioral immobility. N=64 flies/genotype.

### QPCR

Total RNA was extracted from male brains (4 brains/replicate) using the RNAqueous Micro kit (AM1931; Ambion). Manufacturer’s instructions were followed except the lysates were reloaded after the initial filtration, to increase RNA binding, and the RNA was eluted using 75°C RNase-free water (AM9938; Ambion). Following DNase treatment, equal amounts of RNA were reverse-transcribed to cDNA (30-40 ng) using the Superscript III First Strand kit (18080-051; Invitrogen) according to manufacturer’s instructions. QPCR was performed using a 1/10 dilution of the cDNA. The following oligos were used to assess *E74B* and *Cyp18a1* levels in *Drosophila* brains:

mRPS24 Fw: GAACACGTCAACAATGAAGGA

mRPS24 Rv: ACAAACTTGCGGATGAACAC

Aps Fw: GTGTATTCGAGAACAATGACCA

Aps Rv: GTCACGTTCATCACGAACAC

Glo Fw: AATTGGGCACAGGTATATTGAG

Glo Rv: CTCTTCATTTCAGCAATCGAG

E74B Fw: ATCGGCGGCCTACAAGAAG (Caldwell et al., 2005),(Neuman et al., 2014)

E74B Rv: TCGATTGCTTGACAATAGGAATTTC (Caldwell et al., 2005),(Neuman et al., 2014)

Cyp18a1 Fw: 933 AAGAATCACGAGGAGCAACTG

Cyp18a1 Rv: 1033 AGCATGAACACGTTTATCCAC

For each cDNA sample, the data were normalized to the geometric mean of three reference genes (mRPS24, Aps and Glo) as described in Vandesompele *et al*. (2002). The following annealing temperatures were used: 60 °C (mRpS24), 59 °C (E74B), 56 °C (Cyp18a1) and 53 °C (Glo and Aps). Statistical analyses were performed using GraphPad Prism 6 software. At least 3 biological replicates and 3 technical replicates were performed.

### Statistical analyses for *Drosophila* studies

Graphpad Prism software and Microsoft Excel were used for statistical analyses of *Drosophila* fluorometry, immunofluorescence quantification, QPCR and all behavioral assays using ANOVA or unpaired, two-tailed T tests.

### Mice

Mouse lines FVB, 1487 (Mdr1a-/-; Mdr1b-/- (Mdr1)) and 2767 (BCRP^-^/^-^) were purchased from Taconic. Mice bred to maintain homozygosity or purchased were used for non-behavioral experiments. For behavioral experiments, matings starting with FVB and Mdr1 mice were used to generate Mdr1 controls (Mdr1a+/+;Mdr1b+/+ and Mdr1a+/-; Mdr1b+/-) and Mdr1 mutant (Mdr1a-/-; Mdr1b-/-) mice. C57BL/6 mice used as “intruder” were purchased from Simonsen. All mouse protocols were approved by the UCSF and UCSD Institutional Animal Care and Use Committee.

### Collection of mouse brain and blood

Samples were collected from 3.5-month-old males. Mice were anesthetized with ketamine/xylazine/acepromazine maleate mix at 100/6/1mg per Kg mass prior to surgery. Procedures for collecting blood that reduce red blood cell lysis were used. Excess peritoneal fluid was removed through a small opening in the abdomen to prevent flow of fluid into chest cavity, the chest cavity opened, the right atrium cut, and blood pooled in the chest cavity was pipetted with an enlarged-bore tip. Animals were then perfused with cold DPBS to remove blood. Whole brains were dissected, immediately placed in a microcentrifuge tube, frozen in dry ice, and homogenized for analysis or placed at -80°C for later homogenization. Control and mutant mouse samples were collected concurrently.

### Serum preparation for mouse samples

Freshly collected blood was incubated at 37°C for 15 min then at 4°C for 1h to promote clotting. Clotted blood was then centrifuged at 4°C, 5000 RCF for 10 min. Straw-colored supernatant was transferred into a new tube and centrifuged at 4°C, 7500 RCF for 10 min. Supernatant was then transferred to a new tube and stored at -80°C prior to analysis.

### Mass Spectrometry for mouse samples

Targeted metabolites were measured by HPLC tandem mass spectrometry (HPLC-MS/MS) in multiple reaction monitoring mode (MRM) using a SCIEX API 4000 QTrap by Biocrates Life Sciences AG. Seven-point external calibration curves and 13 isotope-labeled internal standards were used. Metabolite concentrations of each sample were determined in a single analysis. Serum samples were directly used for analyses. Brain tissue samples were extracted for steroids using a fixed ratio of extraction volume to weight.

### Pharmacological inhibition of Mdr1

Adult male FVB mice were injected with cyclosporin A (Cell Signaling 9973S; solubilized in 100% DMSO at 50mg/Kg body weight (bw)) or vehicle intraperitoneally at time zero (T=0). All mice were then injected with aldosterone (Sigma A9477; solubilized in 6.25% Ethanol and 6.25% DMSO at 14mg/Kg bw) intravenously by tail vein at T=1hr. The amounts of ethanol and DMSO given to mice were below the LD50 for both chemicals. The following conditions were compared: 1. cyclosporin A vehicle followed by aldosterone, 2. cyclosporin A followed by aldosterone. At T=4hr, brains were collected and analyzed by HPLC-MS/MS as described in sections “Collection of mouse brain and blood” and “Mass Spectrometry for mouse samples.”

### Behavioral tests for mice

Control and mutant male mice used for behavioral tests were born and raised in our animal facility, weaned at age P20, housed with male siblings until social isolation (SI) 3-7 days prior to first behavioral test at 3.5 months of age. Animals were raised with 12:12 hour light:dark cycle. Two sets of behavioral tests were conducted: 1) Open Field (OF)-Elevated Zero Maze (EZM): SI 3 days prior to OF, 2 days SI until EZM. A single cohort of control and mutant mice were tested together. 2) Resident Intruder (RI)-OF-EZM: SI for 7 days prior to RI, 4-5 days SI until OF, 2 days SI until EZM. Two cohorts were tested on different days. Mice were acclimated in the testing room 25-45 minutes before tests. EZM tests were conducted at the UCSF Neurobehavioral Core for Rehabilitation Research during 9am-1pm of the light cycle. Apparatus: 5cm wide circular track of 21.25cm outer diameter, 80cm high with 42cm long open or closed sections, 31 surface infrared beams. Genotypes were blind to operators. Kinder Scientific infrared-detection apparatus and Motor Monitor software were used for behavioral tests. Tests were performed with the operator either in a different room or out of sight from mice being tested, in an activity-absent room with ∼70 decibels of white noise.

### Statistical analyses for mouse studies

Graphpad Prism software was used for statistical analyses of mass spectrometry and animal behavior data using unpaired, nonparametric Mann-Whitney tests.

### Associations between targets and ADRs

We collected 10,098 documented ADRs for 1,332 FDA-approved drugs from the OFFSIDES database (Tatonetti et al., 2012). To avoid duplicates, we standardized and converted each drug to its InChIKey, yielding 1104 unique drug structures. ADRs were considered associated with a drug if OFFSIDES reported a p-value for the drug-ADR link of p < 1×10^-2^; this yielded 9366 unique drug-ADR pairs. We then identified all known human targets for all drugs using ChEMBL (Version 17; (Gaulton et al., 2012)). Targets were considered “known” if a documented ligand for the target had a Tanimoto Coefficient of 1.0 with the drug in question; this yielded 805 (out of 1884 possible) unique targets, 631 of which were human. An initial filter required all drugs to have an association with both an ADR and a target to be considered for analysis (441 drugs, post-filter). In total there were 6,780,515 unique drug-target-ADR triplets. Each unique target-ADR pair had to occur at least 10 times before being considered for enrichment. All resulting drug-ADR (315,279), drug-target (3136), and target-ADR (1,018,208) pairs were then enumerated. An enrichment factor (EF) was calculated for each target-ADR pair after correction for multiple hypothesis testing using the Benjamini-Hochberg method as previously described (Lounkine et al., 2012).

### Alternate target predictions

We used the Similarity Ensemble Approach (Keiser et al., 2007) with RDKit (rdkit.org) ECFP4 (Morgan) and path-based fingerprints to identify predicted additional off-targets of all Mdr1 inhibitors (Table 5) and substrates.

**Table 5:** Top 100 SEA Target Predictions for Mdr1 Compounds. Target predictions for compounds associated with Mdr1. Predictions are generated using the Similarity Ensemble Approach (SEA). All non-human targets are filtered out. Each target is represented as a ChEMBL identifier (ChEMBL 17) uniquely mapped to a Uniprot gene name. If no Uniprot mapping is available, the ChEMBL identifier is used instead. Target frequencies are calculated based on the number of unique compounds predicted to bind with each target. Notably, for the list of Mdr1-associated compounds, Mdr1 is the most frequently occurring target prediction. Max TC represents the maximum Tanimoto Coefficient when measuring similarity between a compound and a target’s ligand set. A Max Tc of 1.0 indicates the compound is a known ligand of the target in question. P-values for each compound-target prediction, of the same target, are averaged and the negative log of the average is calculated. Higher values are indicative of the compound-target interaction being more likely to occur.

| Target | Target Description | Frequency of Predictions | Max Tc | Avg -Log SEA P-value |
| --- | --- | --- | --- | --- |
| MDR1_HUMAN | P-glycoprotein 1 | 13 | 1.00 | 46.78 |
| SCN1A_HUMAN | Sodium channel alpha subunits; brain (Types I, II, III) | 10 | 0.36 | 28.70 |
| HRH1_HUMAN | Histamine H1 receptor | 9 | 0.67 | 36.78 |
| EBP_HUMAN | 3-beta-hydroxysteroid-delta(8),delta(7)-isomerase | 8 | 0.65 | 37.35 |
| DRD5_HUMAN | Dopamine D5 receptor | 8 | 0.67 | 23.61 |
| CCR1_HUMAN | C-C chemokine receptor type 1 | 7 | 0.62 | 81.33 |
| DRD2_HUMAN | Dopamine D2 receptor | 7 | 0.67 | 66.72 |
| CAC1G_HUMAN | Voltage-gated T-type calcium channel alpha-1G subunit | 7 | 0.31 | 34.78 |
| S47A1_HUMAN | Multidrug and toxin extrusion protein 1 | 6 | 1.00 | 22.97 |
| DRD3_HUMAN | Dopamine D3 receptor | 6 | 0.67 | 67.75 |
| ADA1A_HUMAN | Adrenergic receptor alpha-1 | 6 | 0.32 | 33.22 |
| SGMR1_HUMAN | Sigma opioid receptor | 6 | 0.41 | 28.08 |
| 5HT7R_HUMAN | Serotonin 7 (5-HT7) receptor | 6 | 0.46 | 45.54 |
| SC6A4_HUMAN | Serotonin transporter | 6 | 0.42 | 38.05 |
| PLD2_HUMAN | Phospholipase D2 | 5 | 0.90 | 82.25 |
| DRD1_HUMAN | Dopamine D1 receptor | 5 | 0.67 | 23.45 |
| DRD4_HUMAN | Dopamine D4 receptor | 5 | 1.00 | 51.49 |
| ADA2B_HUMAN | Alpha-2b adrenergic receptor | 5 | 0.87 | 23.43 |
| 5HT1A_HUMAN | Serotonin 1a (5-HT1a) receptor | 5 | 0.35 | 51.47 |
| KCNH2_HUMAN | HERG | 5 | 1.00 | 18.30 |
| HRH2_HUMAN | Histamine H2 receptor | 4 | 0.45 | 22.03 |
| SCN2A_HUMAN | Sodium channel protein type II alpha subunit | 4 | 0.31 | 16.53 |
| AOC3_HUMAN | Amine oxidase, copper containing | 4 | 0.44 | 30.37 |
| CP51A_HUMAN | Cytochrome P450 51 | 4 | 1.00 | 82.48 |
| PLD1_HUMAN | Phospholipase D1 | 4 | 0.42 | 114.37 |
| OPRX_HUMAN | Nociceptin receptor | 4 | 0.48 | 42.72 |
| S22A2_HUMAN | Solute carrier family 22 member 2 | 4 | 0.37 | 22.68 |
| HDAC1_HUMAN | Histone deacetylase (HDAC1 and HDAC2) | 4 | 0.30 | 33.05 |
| CCR3_HUMAN | C-C chemokine receptor type 3 | 4 | 0.50 | 24.38 |
| 5HT5A_HUMAN | Serotonin 5a (5-HT5a) receptor | 4 | 0.34 | 19.54 |
| CP24A_HUMAN | Cytochrome P450 24A1 | 4 | 1.00 | 29.06 |
| LKHA4_HUMAN | Leukotriene A4 hydrolase | 4 | 0.31 | 25.32 |
| OPRM_HUMAN | Mu opioid receptor | 4 | 0.38 | 61.91 |
| THB_HUMAN | Thyroid hormone receptor beta-1 | 4 | 0.36 | 28.55 |
| ADRB2_HUMAN | Adrenergic receptor beta | 4 | 0.43 | 99.70 |
| CAN1_HUMAN | Calpain 1 | 4 | 0.39 | 28.59 |
| AMPL_HUMAN | Leucine aminopeptidase | 4 | 0.29 | 16.76 |
| NK2R_HUMAN | Neurokinin 2 receptor | 4 | 0.34 | 54.51 |
| NK3R_HUMAN | Neurokinin 3 receptor | 4 | 0.35 | 30.39 |
| CAC1C_HUMAN | Voltage-gated L-type calcium channel | 4 | 0.35 | 23.12 |
| ADA1B_HUMAN | Alpha-1b adrenergic receptor | 4 | 0.40 | 40.06 |
| CHEMBL2094135 | Gamma-secretase | 4 | 0.32 | 58.59 |
| ADA2C_HUMAN | Alpha-2c adrenergic receptor | 4 | 0.87 | 26.51 |
| ACM3_HUMAN | Muscarinic acetylcholine receptor M3 | 4 | 0.30 | 33.37 |
| ADA1D_HUMAN | Alpha-1d adrenergic receptor | 4 | 0.31 | 30.56 |
| SC6A3_HUMAN | Dopamine transporter | 4 | 0.37 | 14.17 |
| RENI_HUMAN | Renin | 4 | 0.37 | 79.27 |
| IGF1R_HUMAN | Insulin-like growth factor I receptor | 3 | 0.34 | 25.58 |
| CHEMBL2096671 | Serotonin 2 (5-HT2) receptor | 3 | 1.00 | 50.60 |
| ACM1_HUMAN | Muscarinic acetylcholine receptor M1 | 3 | 0.32 | 29.65 |
| CATS_HUMAN | Cathepsin S | 3 | 0.30 | 17.94 |
| ADRB1_HUMAN | Beta-1 adrenergic receptor | 3 | 0.42 | 129.88 |
| CP2J2_HUMAN | Cytochrome P450 2J2 | 3 | 0.36 | 129.04 |
| GRM5_HUMAN | Metabotropic glutamate receptor 5 | 3 | 0.35 | 16.03 |
| ACM5_HUMAN | Muscarinic acetylcholine receptor M5 | 3 | 0.30 | 40.95 |
| CATL1_HUMAN | Cathepsin L | 3 | 0.31 | 16.95 |
| CHEMBL2095200 | Serotonin 2 (5-HT2) receptor | 3 | 0.49 | 25.36 |
| CHEMBL2095166 | Adrenergic receptor beta | 3 | 0.42 | 64.76 |
| CHEMBL612444 | Mitochondrial complex V; ATP synthase | 3 | 0.31 | 50.43 |
| ACM4_HUMAN | Muscarinic acetylcholine receptor M4 | 3 | 0.30 | 37.00 |
| ACM2_HUMAN | Muscarinic acetylcholine receptor M2 | 3 | 0.37 | 27.13 |
| CHLE_HUMAN | Butyrylcholinesterase | 3 | 0.41 | 25.05 |
| ADA2A_HUMAN | Alpha-2a adrenergic receptor | 3 | 0.30 | 26.17 |
| CP17A_HUMAN | Cytochrome P450 17A1 | 3 | 1.00 | 14.32 |
| KIF11_HUMAN | Kinesin-like protein 1 | 3 | 0.32 | 24.02 |
| OPRK_HUMAN | Kappa opioid receptor | 3 | 1.00 | 62.72 |
| OPRD_HUMAN | Delta opioid receptor | 3 | 0.33 | 65.39 |
| MTR1B_HUMAN | Melatonin receptor 1B | 3 | 0.33 | 17.27 |
| BACE1_HUMAN | Beta-secretase 1 | 3 | 0.36 | 35.45 |
| SC6A2_HUMAN | Norepinephrine transporter | 3 | 0.37 | 23.15 |
| ACRO_HUMAN | Acrosin | 3 | 0.31 | 19.26 |
| CATE_HUMAN | Cathepsin E | 3 | 0.34 | 54.20 |
| THA_HUMAN | Thyroid hormone receptor alpha | 3 | 0.36 | 23.46 |
| CHEMBL2094111 | Adrenergic receptor alpha | 3 | 0.39 | 21.15 |
| HYES_HUMAN | Epoxide hydratase | 3 | 0.36 | 56.65 |
| ADRB3_HUMAN | Beta-3 adrenergic receptor | 3 | 0.42 | 54.45 |
| ADRB2_HUMAN | Beta-2 adrenergic receptor | 3 | 0.42 | 108.14 |
| HCN4_HUMAN | Potassium/sodium hyperpolarization-activated cyclic nucleotide-gated channel 4 | 3 | 0.31 | 26.75 |
| CASR_HUMAN | Calcium sensing receptor | 3 | 0.42 | 43.30 |
| CHEMBL2096989 | Angiotensin-converting enzyme | 3 | 0.30 | 27.98 |
| HRH3_HUMAN | Histamine H3 receptor | 3 | 0.67 | 36.93 |
| CATD_HUMAN | Cathepsin D | 3 | 0.41 | 48.91 |
| THAS_HUMAN | Thromboxane-A synthase | 3 | 0.33 | 21.81 |
| MMP8_HUMAN | Matrix metalloproteinase 8 | 2 | 0.36 | 15.54 |
| CHEMBL2096673 | Small conductance calcium-activated potassium channel | 2 | 0.32 | 37.28 |
| CHEMBL2095159 | Serotonin 1 (5-HT1) receptor | 2 | 0.34 | 23.11 |
| CHEMBL2095158 | Adrenergic receptor alpha-2 | 2 | 0.28 | 17.07 |
| CHEMBL2095150 | Phosphodiesterase 1 | 2 | 0.29 | 48.84 |
| FAAH1_HUMAN | Anandamide amidohydrolase | 2 | 0.31 | 20.20 |
| CHEMBL2094116 | Serotonin 3 (5-HT3) receptor | 2 | 1.00 | 24.43 |
| EST1_HUMAN | Acyl coenzyme A:cholesterol acyltransferase | 2 | 0.30 | 17.60 |
| AMPN_HUMAN | Aminopeptidase N | 2 | 0.31 | 18.02 |
| CHEMBL2094125 | Phosphodiesterase 3 | 2 | 0.29 | 32.30 |
| OX2R_HUMAN | Orexin receptor 2 | 2 | 0.37 | 64.24 |
| OX1R_HUMAN | Orexin receptor 1 | 2 | 0.37 | 61.35 |
| PSB2_HUMAN | Proteasome Macropain subunit | 2 | 0.29 | 19.88 |
| MMP2_HUMAN | Matrix metalloproteinase-2 | 2 | 0.80 | 20.90 |
| ANM3_HUMAN | Protein arginine N-methyltransferase 3 | 2 | 0.30 | 14.05 |
| MCHR1_HUMAN | Melanin-concentrating hormone receptor 1 | 2 | 0.64 | 23.29 |
| KCNA3_HUMAN | Voltage-gated potassium channel subunit Kv1.3 | 2 | 0.28 | 16.99 |

## Author contributions

Experiments were conceived by S.J.H., R.N.M., E.D., G.G., H.I., T.K., M.K., M.P.J., R.D., and R.J.B. Experiments were performed by S.J.H., R.N.M., E.D., G.G., S.O., H.I., A.S., and M.D. Data were analyzed by S.J.H., R.N.M., E.D., G.G., H.I., M.K., and A.S. Manuscript was written by S.J.H., R.N.M., E.D., G.G., H.I., T.K., M.K., M.P.J., R.D., and R.J.B.

## Acknowledgements

R.N.M. was funded by UCSF Dept. of Clinical Pharmacology and Therapeutics (GM007546) and Dept. of Anesthesia and Perioperative Care (GM008440) NIH T32 grants. R.J.B. was supported by NIH GRANTs R01GM081863 and R21NS082856, NIEHS R21 ES021412, the UCSF Department of Anesthesia and Perioperative Care, and the Sandler foundation. R.D. was funded by the UCSF Program for Breakthrough Biomedical Research. M.J.K. was funded by a NIH SBIR (GM093456) and a Glenn Foundation Award for Research in Biological Mechanisms of Aging. G.G. was supported by iPQB Bioinformatics Training Grant. We would like to acknowledge members of the Kenyon and O’Farrell labs for critical reading of the manuscript and helpful suggestions. We would also like to thank members of the Bainton, Daneman and Kroetz labs for technical assistance and useful guidance during this work.

### Conflict of Interest Statement

M.P.J. is a consultant to Schrodinger LLC, which licenses, develops, and distributes software used in this work.

